# Intra-specific evidence to support the drift-barrier hypothesis of mutation rate evolution

**DOI:** 10.64898/2026.07.31.741990

**Authors:** Chaowei Zhang, Hongbo Wang, Kerry Reid, Dandan Wang, Lei Lv, Yvette Heimbrand, Mikkel Heide Schierup, Juha Merilä

## Abstract

Rates of *de novo* mutations (*µ*) are highly variable among different taxa but less is known about their variability at the intraspecific level. Using a large data set (n=364 trios) from eight nine-spined stickleback (*Pungitius pungitiu*s) populations differing in their effective population sizes (*N_e_*), we tested the prediction of the ‘drift-barrier hypothesis’ (DBH) that *µ* scales negatively with increasing *N_e_*. Indeed, *µ* was a negative function of *N_e_*, also after correcting for phylogenetic non-independence of populations. While the range of variation in mutation rates across populations spanned more than a 4-fold range (*µ* = 1.60 - 6.99 × 10^-9^), we discovered one highly mutable family with a five-times’ higher mutation rate than the population average. Evidence was also found (i) for reduced efficiency of selection in small freshwater populations subject to strong genetic drift, (ii) that maternal age associates positively with *µ*, and shorter generation time elevates per-year mutation rates, (iii) that both replication errors and DNA repair efficiency contributed to *µ*, and that (iv) mutation rate variation has a polygenic basis. In general, the results provide support for the DBH identifying elevated mutation rates in populations subject to strong genetic drift.

## Introduction

Germline mutations are the fundamental source of genetic variation. Despite the relatively low per-nucleotide mutation rates (*µ*), they exhibit substantial variability across species (Bergeron et al. 2023; Wang and Obbard 2023). At the molecular level, this variation is governed by the contributions of DNA replication errors and the accumulation of unrepaired damage (Zhu et al. 2025). However, disentangling their relative contributions is challenging due to the complex interplay of multiple factors. For instance, male-biased mutation rates observed in mammals and birds can be a result of the higher number of germline cell divisions, and hence, replication errors, in males compared to females (Bergeron et al. 2023). On the other hand, the positive effect of maternal age on germline mutation rates, despite the absence of cell divisions after puberty, has also been attributed to unrepaired DNA damage (Gao et al. 2019; Wu et al. 2020; Spisak et al. 2024).

While it is hard to distinguish the replication- and DNA-damage derived mutations (Lewin and Eyre-Walker 2025), the drift-barrier hypothesis (DBH) offers a compelling explanation to the observed variation in mutation rates (Sung et al. 2012; Lynch et al. 2016). It suggests that the extent of molecular or cellular perfection, including replication fidelity and DNA repair effectiveness, is modulated by genetic drift, which limits the efficiency of natural selection in purging deleterious mutations or, in rare cases, fixing beneficial ones (Lynch 2010; Lynch 2011; Sung et al. 2012; Lynch et al. 2016). In line with this, the per-generation *µ* decreases as the effective population size (*N_e_*) increases across the tree of life (Lynch 2010; Lynch 2011; Sung et al. 2012). Recent expansions of the DBH highlight that life-history traits like generation-time (cf, the age at sexual maturity) affect the fitness of mutation modifiers (Lewin and Eyre-Walker 2025; Zhu et al. 2025). These studies suggest that the per-year *µ* correlates negatively with generation time, implying that the long-lived species may be selected for more effective damage-repair systems to minimise the lifetime mutation loads (Zhu et al. 2025).

While these explanations are compelling, in addition to *N_e_* and generation time, many other factors likely vary across the species and could affect the observed relationships (e.g., genomic background and structure, reproductive strategy, etc.). Therefore, it would be of interest to see if this relationship holds true also in intraspecific population comparisons in which other possible sources of variation can be better controlled for. However, studies on intraspecific variation in germline mutation rates are rare and typically not well replicated at the population level (Narasimhan et al. 2017; Wang et al. 2023; López-Cortegano et al. 2024; Garcia-Salinas et al. 2025; Zhang et al. 2025), limiting our understanding of what ultimately determines the large differences in mutation rates across the tree of life (Lynch 2010; Lynch et al. 2016; Lewin and Eyre-Walker 2025; Zhu et al. 2025).

Nine-spined sticklebacks (*Pungitius pungitius*) provide an ideal system to study these hypotheses at the intraspecific level. Firstly, *P. pungitius* exhibits a wide range of neutral genetic diversities and levels of inbreeding leading to substantial differences in *N_e_*s (Kivikoski et al. 2023; Chen et al. 2025; Feng et al. 2025). Therefore, efficiency of natural selection in independently evolving freshwater isolates is limited by strong genetic drift, whereas selection is expected to be more efficient in large outbred marine populations (Wang et al. 2026a). Secondly, *P. pungitius* ecotypes differ in generation times by approximately one year (DeFaveri et al. 2014). For a short-lived species, this is a large relative shift providing an opportunity to test the generation time effect on mutation rates, which isolates its role from the massive confounding differences in metabolism and other life-history traits that complicate cross-species studies (Weinstein and Roy 2026).. Additionally, the recurrent mutations among siblings, which are known to originate early during zygotic fertilization (Biesecker and Spinner 2013), can be investigated due to the large number of offspring the species produce. Moreover, in contrast to mammals, female and male fish generate a similar number of germ cells subject to more equal number of cell divisions, making this system particularly interesting to address questions about the etiology of *de novo* mutations, as well as sex bias in mutation rates.

Recent technological advancements have substantially enhanced genome assembly methodologies. The reference genome of *P. pungitius* has been repeatedly and substantially refined, with each update demonstrating significant improvements in quality and contiguity (Varadharajan et al. 2019; Kivikoski et al. 2021; Wang et al. 2024). Although these assemblies were considered to be of high-quality, with version 7 being used extensively (Zhang et al. 2023; Feng et al. 2024; Chen et al, 2025), the phasing of the two sex chromosomes (improved from v7 to v8; Kivikoski et al. 2021; Wang et al. 2024), chrX and chrY, remains incomplete, primarily due to challenges in assembling repetitive regions. In recent years, the integration of highly accurate long-read sequencing (Pacbio HiFi), ultra-long Oxford Nanopore Technologies (ONT) sequencing, and chromosome conformation capture (Hi-C) sequencing has revolutionized genome assembly techniques, enabling the generation of haplotype-resolved telomere-to-telomere (T2T) assemblies (Li and Durbin 2024). Error-eliminated genome assemblies facilitate more accurate identification of mutations across the genome and yield improved estimates of mutation rates, potentially advancing our understanding of the relative importance of different drivers behind mutation rate variation.

The primary aim of this study was to provide an intraspecific test of the drift-barrier hypothesis by asking whether the germline mutation rates scale negatively with *N_e_* across nine-spined stickleback populations, while controlling for the impact from generation time and metabolic drivers. We also tested whether decreased efficiency of selection is associated with increased mutation rates across the study populations by comparing the neutrality index in different populations. The point germline mutation rates were estimated from 364 parent-offspring trios originating from four isolated freshwater and four outbred marine populations by mapping the mutations to a new high-quality T2T reference genome assembly. In an attempt to differentiate between replication errors and DNA damage as the ultimate source of new mutations, we further characterized and compared the mutation spectra across populations, as well as estimated the effects of age and sex on mutation rates.

## Results

### The telomere-to-telomere genome assembly

We successfully constructed a telomere-to-telomere (T2T) genome assembly (*NSPV9_T2T*, NCBI: PRJNA1334546) for a male *P. pungitius* individual from a Baltic Sea population (Tvärminne; TVA, 59.83°N, 23.00°E), utilising 173.8x PacBio HiFi, 84.6x ultra-long ONT, and 389.0x Hi-C data (Table S1, Figure 1A). The initial assembly produced from Hifiasm (0.20.0-r639; Cheng et al. 2024) contained 174 contigs for haplotype 1 and 106 contigs for haplotype 2. We then anchored and oriented the contigs into 42 chromosomes using Hi-C data, and all autosomes were named according to the most recent enhanced genome assembly (version 8; ENA accession ID: GCA_902500615; Wang et al. 2024). Sequencing gaps, especially those involving segmental duplications (SDs), were filled with ultra-long ONT sequences.

**Figure 1.**
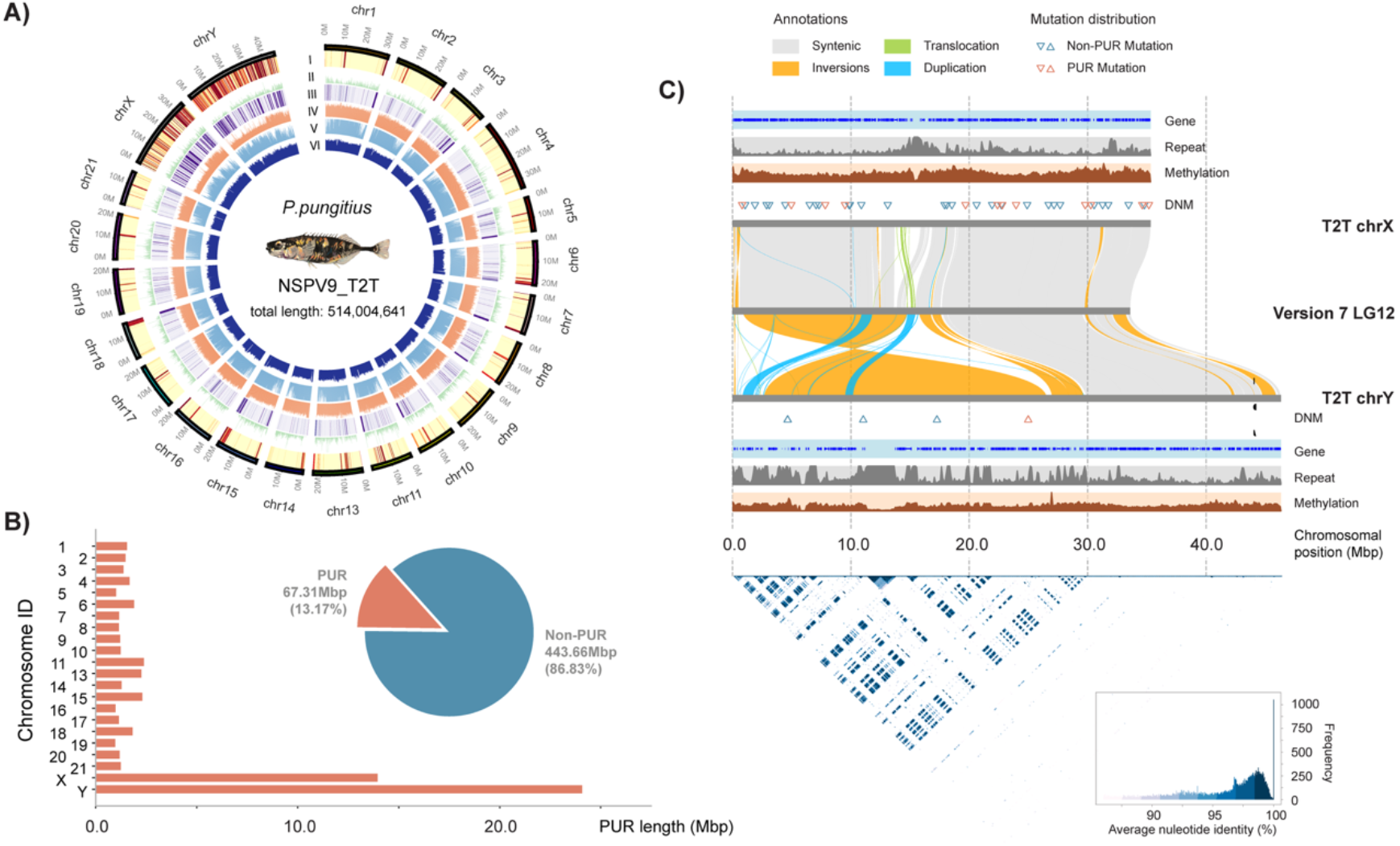
Summary of the complete telomere-to-telomere (T2T) genome assembly. A) Basic information of the improved T2T assembly. The bands demonstrate, sequentially from I to VI, previously unassembled regions in comparison to V7 (red bars), gene density, segmental duplication density, the coverage of ultralong ONT reads, the coverage of HiFi reads, and the methylation level based on HiFi data, in 10-kb windows (Tools involved to generate this graph: plotsr (v1.1)(Goel and Schneeberger 2022); ModDotPlot (v0.9.8)(Sweeten et al. 2024). B) The per-chromosome length and whole genome proportion of the previously unassembled regions (PUR). C) Comparison of the chrX and chrY in the T2T assembly with the unphased sex chromosome in the Version 7 (LG12). The identified DNMs are marked with triangles for the two sex chromosomes. The heatmap at the bottom displays the pairwise average nucleotide identity (ANI) between homologous sequences in the sex determination region of chrX and chrY, with annotations of the color scales at the bottom right corner.

*NSPV9_T2T* enhanced the assembly quality of sex chromosomes (chrX and chrY) in particular. The hybrid assembly approach of Hifiasm and Verkko was employed to address the challenges brought by the longer length of chrY compared to chrX. The local quality value (QV) and mapping rate for chrY produced by Hifiasm surpassed those by Verkko, but it fragmented the SDR region into five contigs. Therefore, we manually anchored and ordered these five contigs based on Verkko’s assembly order, as its assembly exhibited better completeness (Figure 1A). Ultra-long ONT reads were applied for gap filling, and it helped to resolve the highly repetitive regions at 6Mb and 11Mb on chrY. Altogether, these steps constructed a full-length Y chromosome for nine-spined stickleback, with higher contiguousness and completeness compared to previous assembly versions (Figure 1C, Figure S3).

The final assembly consisted of 20 autosomes and chrY from haplotype 1, chrX from haplotype 2, and the mitochondrial genome, yielding a total genome length of 514,004,641 bp (Table 1, Figure S2, S3). It exhibited an equivalent BUSCO (Benchmarking Universal Single-Copy Orthologs) value at 99.0% and a higher QV of 53.3 (base accuracy >99.999%) compared to the version 8 (Table S2). Based on the NCBI Eukaryotic Genome Annotation Pipeline, we annotated 23,760 genes with RNA-seq data from 48 samples (Wang et al. 2020) and high-quality Iso-Seq data from one UK individual (PRJEB59309), including 22,963 protein-coding genes (Table S3, S4). These contained 99.1% *Actinopterygii-*conserved genes in the BUSCO analysis, suggesting the high completeness of the annotation. 114.11 Mb repetitive elements were detected in our assembly, which was 27.41 Mb longer than in the previous version (Table 1). Compared to V7, a total length of 67.31 Mbp previously unassembled region (PUR) was assembled in *NSPV9_T2T* (Figure 1B). These regions included mainly sex chromosomal elements (38.03 Mbp in total) as well as centromeric microsatellites and segmental duplications on the short arms of autosomes ranging from 4.68 to 18.90 Mbp in total length (Figure 1B).

**Table 1.** Comparison of three *Pungitius pungitius* genome assemblies.

| Feature | PYO_v7 | PYO_v8 | NSPV9_T2T |
| --- | --- | --- | --- |
| Genome Size (bp) | 466,582,808 | 466,451,302 | 514,004,641 |
| Chromosome Genome Size (bp) | 439,721,235 | 463,473,258 | 514,004,641 |
| N50 contig size (bp) | 2,775,277 | 1,233,545 | 21,891,256 |
| N50 scaffold size (bp) | 20,450,314 | 20,475,552 | 21,891,256 |
| Y chromosome length (bp) | NA | 22,806,223 | 46,410,677 |
| Total number of contigs | 2,487 | 3,822 | 35 |
| Number of contigs in Y Chromosome | NA | 400 | 6 |
| Number of Gaps | 341 | 527 | 0 |
| BUSCO completeness | 99.1% | 99.2% | 99.0% |
| Quality value (QV) | NA | 35.30 | 53.3 |
| Repeat content | NA | 86.75Mb (18.6%) | 114.11Mb (22.2%) |
| Annotated BUSCO genes | NA | 92.50% | 99.10% |

### Accurate DNM identification utilising the NSPV9_T2T

With the improved reference genome, a total of 935 unique point *de novo* mutations (DNMs) appearing in offspring were identified in 364 trios after manual curation, with 472 from 146 freshwater (FW) and 463 from 218 marine (MA) trios (Table S5). Among these, 890 candidates were located on autosomes, 20 in pseudoautosomal region (PAR), and 25 in sex-determination region (SDR), among which 135 (14.44%) were found on the repetitive regions of the genome. Interestingly, the new genome assembly provided 22 DNMs that were located on the abovementioned PURs of version 7 assembly, 59.09% of them on the sex chromosomes, whereas all autosomal ones were located on the repetitive regions (Figure 2C). The same dataset and pipeline were employed to call DNMs utilising V7 assembly, and a total of 982 DNMs on autosomes and pseudoautosomes were detected. 141 (14.36%) among them were within the repetitive regions. These did not include the SDR region due to its assembly incompleteness (Figure 1C).

**Figure 2.**
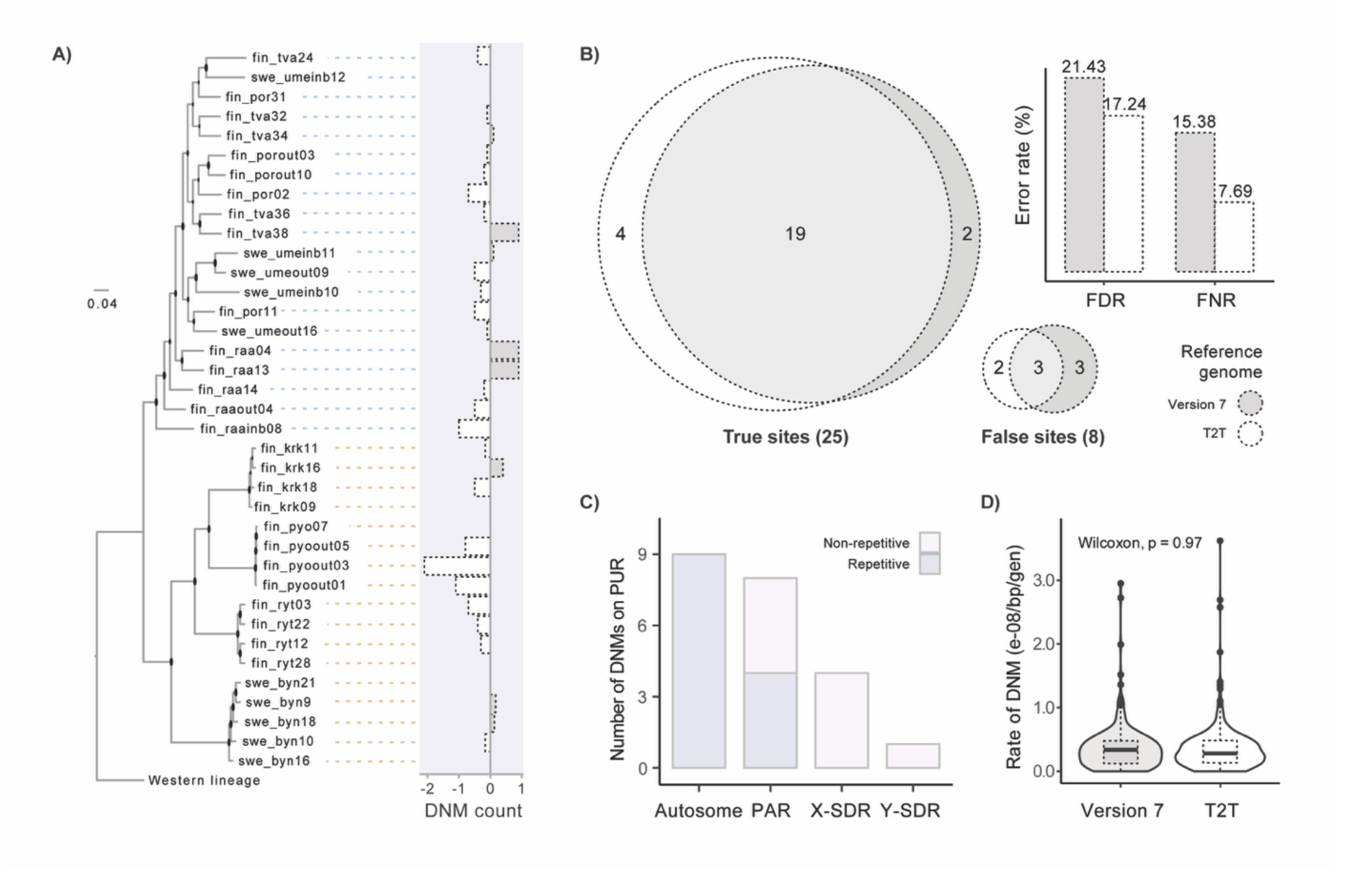
Comparison of the *de novo* mutation (DNM) rates estimated from the two reference genomes. A) The differences in DNM counts utilising the two reference assemblies across pedigrees, with bars on the left representing more DNMs using V7, and bars on the right showing the opposite. B) The Venn plots summarise the results of Sanger sequencing to verify the candidate DNMs identified using the two reference genomes. The bar plot shows the resulting false discovery rates (FDR) and false negative rates (FNR). C) The number of DNM candidates identified on the PUR regions (light grey and light blue: non-repetitive and repetitive regions). D) The final germline DNMs rates for the two assemblies after correcting for FDR and FNR.

To validate the single-nucleotide DNMs called from both assemblies, we performed Sanger sequencing on 350–550 bp flanking regions of 38 candidates from 15 trios. 33 regions were successfully amplified, and 78.8% of them were found to be true DNMs. *NSPV9_T2T* outperformed the V7 reference by presenting both lower false discovery rates (FDR: 17.24% vs. 21.43%) and lower false negative rates (FNR: 7.69% vs. 15.38%; Figure 2B). While the FNR shown in Figure 2B was based on the missed true DNMs detected with the other genome, the filter-FNR (fFNR), accounting for errors introduced by the allelic balance filter, was used in the final formula, averaging at 6.07% and 6.32% for *NSPV9_T2T* and V7. The callable genome sizes (CGS) for the autosomal regions (including PAR) were 345.05 (*NSPV9_T2T*) and 347.79 (V7) Mbp on average, representing 77.41% and 82.26% of these regions in each genome, respectively. Although the pedigree-level DNM counts were generally comparable or sometimes elevated when using the V7 due to its increased error rates (Figure 2A), the mean germline DNM rates on autosomes were almost equal (3.546 and 3.552 × 10^-9^/bp/gen; p=0.97, Figure 2D) after adjusting by fFNR, FDR and CGS (population-level statistics see Table S5).

### Intraspecific variation in de novo mutation rates

The autosomal DNM rates estimated by *NSPV9_T2T* averaged 4.63 and 2.74 × 10^-9^/bp/gen for the FW and MA populations (Table S6), values significantly different at both the pedigree (*t*_28.1_=2.50, p=0.019; Figure 3A) and the individual level (*t*_223_=3.39, p=8.4e- 04; Figure S5A) analyses. The harmonic mean of long-term effective population sizes (*N_e_*) was estimated up to 8.15-fold higher in MA than in FW populations (Table S6, S7). Accounting for the phylogenetic non-independence of different populations by considering their coancestry, a significant decline in per-generation DNM rates with increasing coalescent *N_e_* (log-scaled) was observed (phylogenetic generalised mixed linear model (PGLMM), quasipoisson distribution in MCMCglmm, β=-0.40, p=0.0075, Figure S6A, Table S7A), aligning with the general trend observed interspecifically in other vertebrates (generalised mixed linear model (GLMM) in glmmTMB, β=-0.09, p=0.023, Figure 3B, Table S7B). However, this trend was not significant if contemporary *N_e_* was used (PGLMM, β=-0.09, p=0.070, Table S7, Figure S6B).

**Figure 3.**
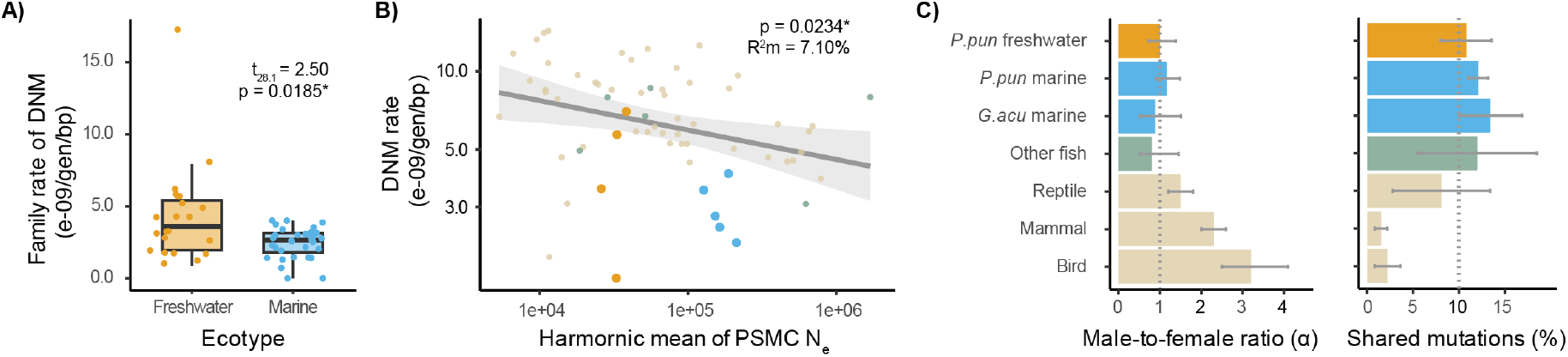
Variation in per-generation *de novo* mutation (DNM) rates. A) The mean pedigree-level DNM rates in the two ecotypes, t-test statistics excluding the hypermutable outlier pedigree. B) Correlation between per-generation DNM rate and the harmonic mean of PSMC *N_e_*. The statistical test results come from glmmTMB Gaussian models. Points in ivory (mammals, birds and reptiles) and green (fish) are from Bergeron et al.(Bergeron et al. 2023); blue: marine populations from this study and Zhang et al.(Zhang et al. 2025) (*Gasterosteus aculeatus*); yellow: freshwater populations. C) Male-to-female ratio (α, Left) and the percentage of shared mutations (right) in different taxonomic groups (*P. pun* and *G.acu* refer to nine- and three-spined sticklebacks, respectively)

To see if there is evidence for decreased efficiency of selection in small freshwater populations, we utilised the neutrality index (*Ni*) derived from the McDonald–Kreitman test (detailed in Materials and Methods) to compare the strength of selection between the ecotypes. While the strength of negative selection varied among genes and populations (Table S6; Figure S8A), the overall strength of selection was stronger in marine than freshwater populations (Table S6, GLMM, p=0.035). The per-generation DNM rates were negatively associated with *Ni* (GLMM, β=-4.444, p=2.89e-09), although this association was not significant after accounting for the phylogenetic relatedness (PGLMM, β=-0.121, p=0.879; Figure S8B). To further validate this, we used the slope of models relating nucleotide diversity to the recombination rate for each population (Table S6) as a proxy of linked selection efficacy (Wang et al. 2026a). In agreement with what was seen for *Ni*, the selection in freshwater populations was more restricted by genetic drift, presenting lower slopes. Correspondingly, the per-generation mutation rate correlates negatively with this slope (GLMM, β=-0.709, p<2e-16), although it is again not so obvious when corrected by phylogeny (PGLMM, β=-0.201, p=0.119).

### Effect of parental age and generation time to germline mutation rates

55.08% and 15.25% mutations were phased back to their parent-of-origins for individuals from MA and FW populations. The α values were low for both ecotypes (FW: 1.0, 95% confidence interval [CI]: 0.7 – 1.4; MA: 1.2, CI: 0.9 – 1.5) and also low when compared to the values from other taxonomic groups (Figure 3C). In contrast, both ecotypes demonstrated an inflated proportion of shared DNMs at 12.1% (MA, se=1.1%) and 10.8% (FW, se=2.8%). These values are similar to those seen in other fish species, including the sister species, the three-spined stickleback (*Gasterosteus aculeatus*, Figure 3C), but with less sex bias (lower α values) and more mutations occurring at the early prezygotic stage (higher proportion of shared DNMs) than in other vertebrates (Figure 3C).

The parental fish aged one to four years old in our dataset, and the FW fish were significantly older than MA fish (*t*_70.8_=7.54, p<0.001). Because of the low proportion of shared mutations and low number of older parents, we did not manage to phase any of the shared DNMs back to the parents older than two years’ old, and therefore, were not able to see any variation in rates of shared mutations associated with parental age. However, after correcting for the phasing ratio, the per-generation rate of non-shared DNMs was significantly higher in older mothers (GLMM, β=0.364, p=0.0084), but not so in the fathers (GLMM, β=-0.021, p=0.910, Figure 4A). The bivariate model, accounting for the joint effect of the maternal and paternal ages, confirmed the positive impact of maternal age to per-generation DNM rates (GLMM, β=0.536, p=1.58e-03), but again, not for the paternal age (GLMM, β=0.222, p=0.365).

**Figure 4.**
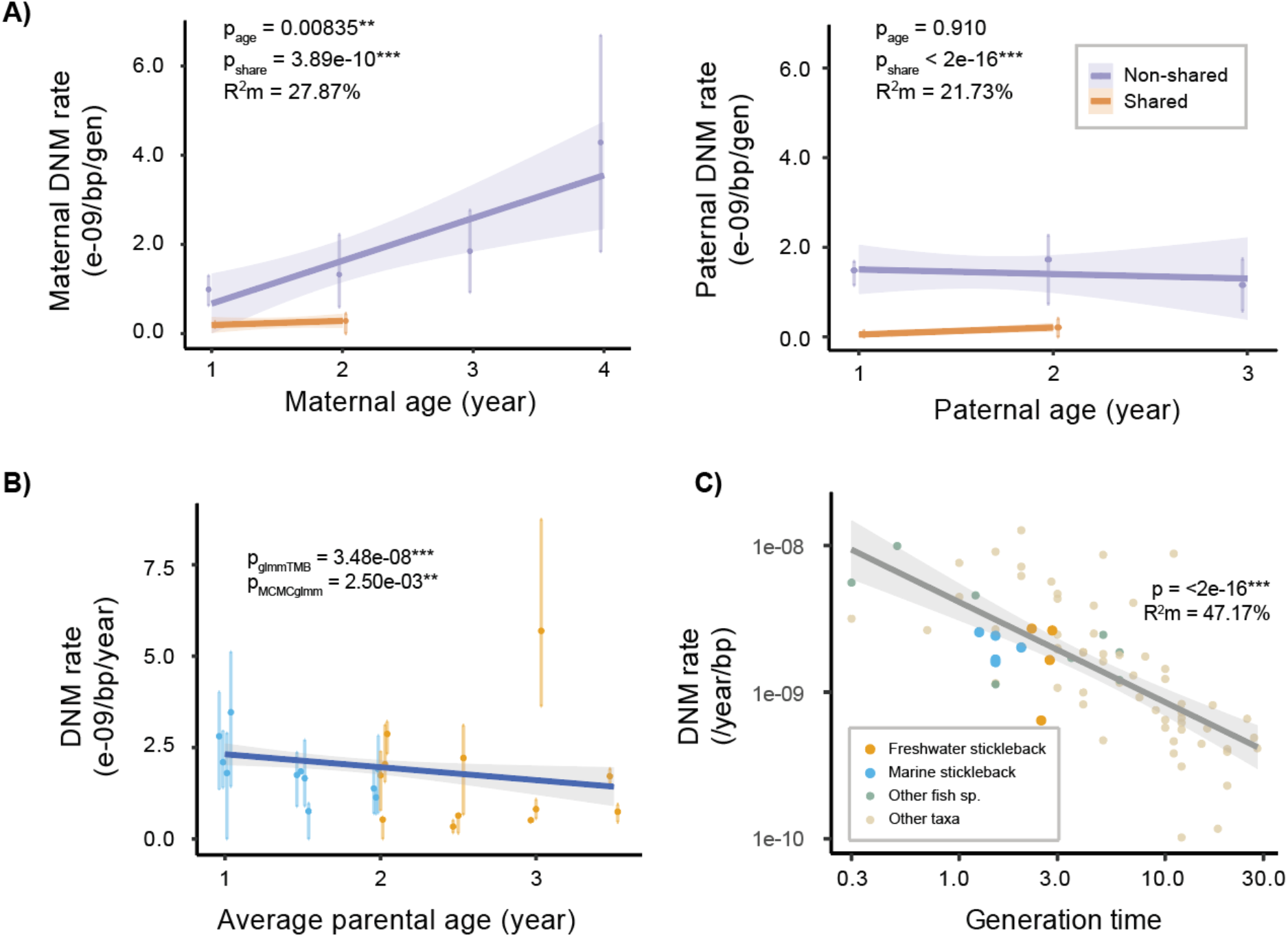
Effect of parental age and generation time to germline mutation rates. A) Association of shared (orange) and non-shared (purple) DNM rates (/gen/bp) with maternal (left) and paternal (right) age. B) Association between the per-year DNM rate and average parental age in nine-spined sticklebacks (blue: marine pedigrees, yellow: freshwater pedigrees). C) Association between per-year DNM rate and the generation time. The statistical test results come from glmmTMB Gaussian models. Points in ivory (mammals, birds and reptiles) and green (fish) are from Bergeron et al.(Bergeron et al. 2023); blue: marine populations from this study and Zhang et al.(Zhang et al. 2025) (*Gasterosteus aculeatus*); yellow: freshwater populations

Dividing the per-generation DNM rate by the mean parental age, the estimated per-year DNM rate was 2.04 × 10^-9^/bp/yr on average. We found evidence for significant negative correlation between yearly DNM rate and parental age (GLMM, β=-0.46, p=3.48e-08; PGLMM, β=-0.49, p=2.50e-03; Figure 4B) and the population generation time (GLMM, β=-0.25, p=4.56e-03; PGLMM, β=-0.27, p=0.428) when excluding the hypermutable outliers, consistent with the interspecific trend (Figure 4C). The per-year DNM rate also displayed a subtle distinction between the two ecotypes, with the FW ecotype displaying an average rate 7.11e-11(/year/bp) lower than the MA ecotype (t_327_=-2.05, p=0.041; Figure S5B). Converse to the trend for per-generation DNM rates, the yearly DNM rate correlated positively with the coalescent *N_e_* (GLMM, β=0.430, p=5.52e-10, Figure S7), suggesting a strong impact of the generation time on the realised mutation accumulation per unit of time.

### Mutation spectra and the hypermutable pedigree

Significantly more DNMs occurred on C:G sites than on A:T sites given the nucleotide constitution of the genome (*χ²* = 275.89, df = 1, p < 0.001), with a larger proportion of C>T mutations (46.52%) among which 51.95% were methylated CG dinucleotides (CpG > TpG, Figure 5B). On average 16.47% and 14.12% involved C>A and T>C mutations, ranking sequentially the second and the third common types. Based on these, 60.64% and 39.36% DNMs were transitions (Ts) and transversions (Tv), respectively, showing a Ts:Tv ratio at 1.54 for all samples. This ratio was not different for the two ecotypes (FW: 1.65 vs. MA: 1.44; t_39.5_=1.15, p=0.256; Figure 5C, Figure S9A). Additionally, the proportion of strong-to-weak (S>W) DNMs was slightly higher than that of weak-to-strong (W>S) notably in FW populations, resulting in an increased S>W/W>S ratio although it was similar in both ecotypes (FW: 3.77 vs. MA: 3.05; t_41_=1.14, p=0.26; Figure 5C, Figure S9B). Compared to MA individuals, the mutation spectra were more variable in FW ones, exhibiting wider confidence intervals for both Ts:Tv and S>W/W>S ratios (Figure 5C).

**Figure 5.**
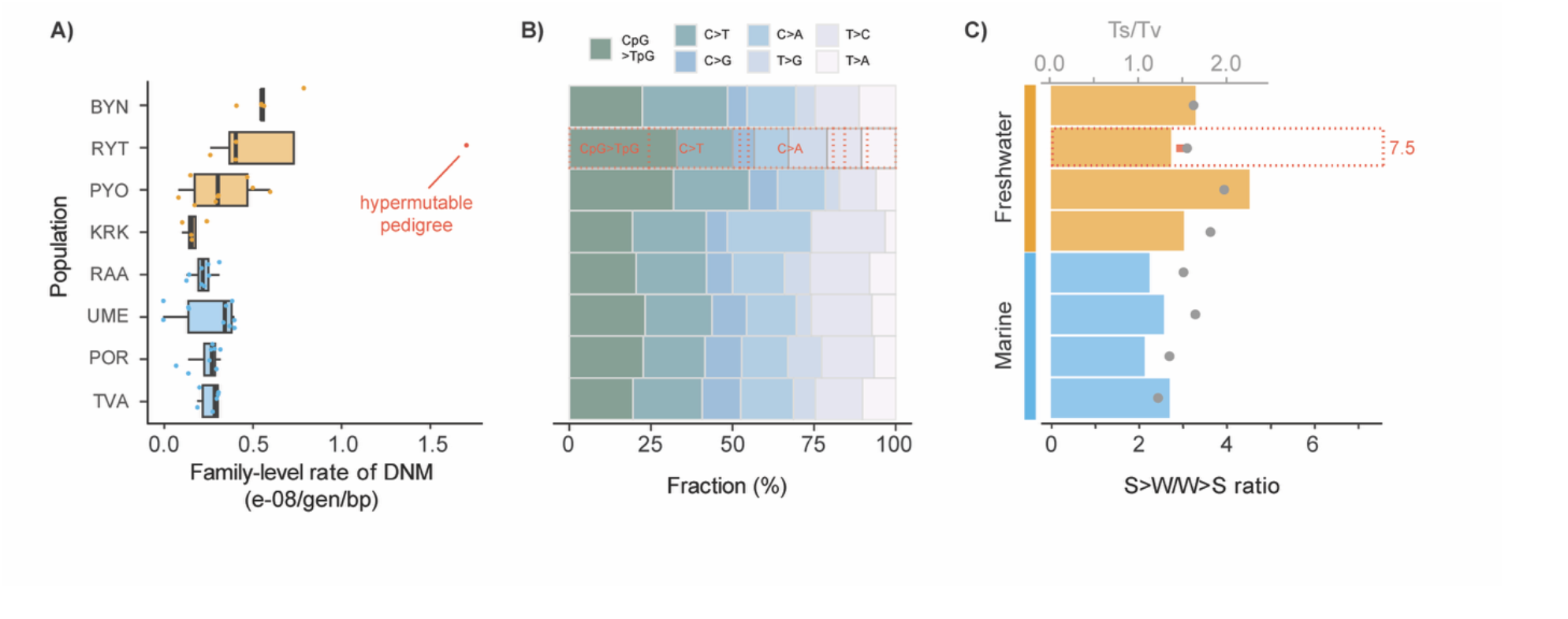
The mutation spectrum in different stickleback populations 1079 including the hypermutable pedigree. A) The pedigree-level germline de novo mutation rates in eight populations, with the hypermutable pedigree, fin_ryt22, marked in red. B) The mutation spectra for the eight populations (fin_ryt22 in red dotted outlines). C) The ratio of strong-to-weak (S>W)/weak-to-strong (W>S) mutations in the eight populations (bars, lower primary x-axis), and the transition (Ts) over transversion (Tv) ratio (points, upper secondary x-axis). fin_ryt22 demonstrated a much higher S>W/W>S ratio (red dotted1086outline bar) but similar Ts/Tv value compared to the other RYT pedigrees (red dot).

An exceptionally hypermutable pedigree (*fin_ryt22*) was found in one of the freshwater populations (Rytilampi; RYT, 66.38°N, 29.32°E), whose mutations occurred at a five-times’ higher rate (1.71 × 10^-8^/bp/gen) than the average DNM rate (Figure 5A). This pedigree also had significantly higher DNM rates than the other pedigrees of the same or even older average parental age (>3 yrs; t_9.35_=3.88, p=0.003). This pedigree demonstrated a unique mutation spectra as compared to the other pedigrees of the same population with more C>A mutations (Figure 5B, χ² = 48.04, df = 6, p = 1.16e-08), presenting a significantly higher S>W/W>S ratio at 7.5 (χ² = 14.06, df = 1, p = 1.77e-04), whereas the Ts/Tv ratio was similar to that in other pedigrees (Figure 5C, χ² = 0.09, df = 1, p = 0.76). Although not appearing in all offspring, a loss-of-function (LOF) mutation was found on a DNA-repair related gene, *ANKLE*, in one of the male offspring. It was a stop-gained CpG>TpG mutation transmitted from his father.

Finally, genome wide association study (GWAS) identified 50 significant quantitative trait loci (QTL) to be associated with DNM rates, and the gene ontology enrichment analyses indicated that neuron fate commitment, animal organ formation and cell adhesion pathways were the most associated (Figure S10). Among these SNPs, two demonstrated uniquely in the both parents of *fin_ryt22*, one of which was caused possibly by a deletion (chr10: 4,393,854-4,393,920; Figure S11) located on an uncharacterised long non-coding RNA between a gap junction gene, *gja4* (chr10: 4,449,446 - 4,452,838) and a signalling transmitter, *dlgap3* (chr10: 4,355,142 - 4,366,055), whereas another was located in the intronic region of *myo6a* gene.

## Discussion

To the best of our knowledge, this is the first study providing an intraspecific test of the drift-barrier hypothesis (DBH). As predicted by the hypothesis, we observed a negative correlation between per-generation point germline *de novo* mutation (DNM) rates and effective population sizes (*N_e_*). Further in support of the premises of DBH, we found evidence for reduced strength of selection in small freshwater populations subject to strong genetic drift. Contrary to this trend, the per-year DNM rate increased with increasing *N_e_*, and decreased with the increasing generation time, suggesting that generation time plays an important role in the DBH context. In fact, we found a positive relationship between maternal (but not paternal) age and mutation rate, suggesting that older mothers contribute disproportionately to mutation rates. However, female and male mutation rates were very similar, likely due to similar numbers of cell divisions between the two sexes in fish (Bergeron et al. 2023). Interestingly, we also observed a hypermutable family in one of the low *N_e_* populations, likely resulting from elevated DNA damage. The new high-quality telomere-to-telomere reference genome allowed us to identify nearly a thousand unique DNMs, but after correction with a subset of DNMs validated through Sanger sequencing, 17% of these were deemed to be false positives underscoring the difficulty of calling DNMs solely based on bioinformatic data. In the following, we will discuss these findings in detail and explore their implications for our understanding of mutation rate variation.

The drift-barrier hypothesis predicts a negative relationship between mutation rate and long-term *N_e_*, and this was what we observed across the eight nine-spined stickleback populations. The per-generation mutation rates in smaller freshwater (FW) populations were higher than those in larger outbred marine populations. That this relationship was not significant when using current *N_e_* is not surprising as the contemporary *N_e_* may not reflect the effective size of these populations when the mutation rates were evolving. Interestingly, FW populations also exhibited greater variability in mutation rates, even more than what is observed between them and marine (MA) populations. Among them, three pond populations located close to each other in northeastern Finland, possibly sharing similar exogenous (e.g. temperature, conductivity, etc), and endogenous conditions (e.g. metabolic rate, cell division rate, etc.), display markedly different mutation rates and spectra. This variation could stem from differences in unrepaired DNA damage, partly due to repair efficiency, but also from differences in stress they have experienced considering the possibility of high environmental stress associated with external fertilisation in this species. Both GWAS results (highlighting roles for DNA repair, cell communication, and stress pathways) and the hypermutable pedigree (notably involving the *ANKEL* gene linked to DNA repair) underscore the combined importance of the DNA damage and repair efficiency in sticklebacks. Furthermore, the unique mutation spectrum with higher proportion of C>A mutations that has been consistently observed in different populations and species of sticklebacks (Zhang et al. 2023; Zhang et al. 2025), other fish species (Beichman et al. 2023; Bergeron et al. 2023), and mollusks (Wooldridge et al. 2025) is suggestive that the unrepaired environment-induced DNA damage is a primary cause of *de novo* mutations.

The effectiveness of these repair and defense mechanisms is shaped by varying levels of natural selection constrained by genetic drift, as proposed by the drift-barrier hypothesis (Lynch 2010; Lynch 2011; Sung et al. 2012). While it is reasonable to assume that the external sources of mutagenesis should be fairly similar across different pond populations, there may be population differences in internal stress tolerance and DNA repair efficiency. For instance, we know that the levels of inbreeding and loads of deleterious mutations vary among these populations (Chen et al. 2025) and this variation could contribute to the observed variation in mutation rates. It is also interesting to note that the pond population with the lowest mutation rate (KRK) differs from other studied pond populations in that it has experienced a bout of piscine predation in the recent past due to the introduction but now extirpated brown trout (Herczeg et al. 2010; Fraimout et al. 2022). One can speculate that brown trout predation may have removed individuals carrying many *de novo* mutations when population size crashed, and genetic diversity decreased.

The results demonstrate a clear interplay between the population-level selection and the individual-level parental effects on mutation rates. While older parents transmitted more mutations to their offspring, the rate of this accumulation is not uniform among populations. Although *N_e_* and breeding age are often colinear, freshwater parents exhibit significantly higher mutation rates than those of marine parents of the same age (age = 2, t_165_ = 2.43, p = 0.01). In marine populations with larger *N_e_* and higher selection efficiency, the fitness cost of the increased mutation load associated with older parental age is penalized. However, for small freshwater populations, the less efficient selection limited by stochastic genetic drift may prevent the refinement of DNA repair modifiers.

While recent cross-vertebrate studies suggest that long-lived species have evolved more effective DNA repair systems to maintain low yearly mutation rates reducing the lifetime mutation load (Zhu et al. 2025), our intraspecific data demonstrate a different dynamic. Our observation that yearly DNM rate is lower in long-lived freshwater ecotypes, despite their inferior molecular fidelity at age 2, suggests a ‘buffering’ effect of generation time. This is consistent with findings in other vertebrates where a substantial ‘intercept’ of mutations is acquired during early developmental bursts of cell division (Gao et al. 2019; De Manuel et al. 2022; Lewin and Eyre-Walker 2025). The results further demonstrate that the ‘molecular clock’ can be inconsistent within a single species. However, we consider these observations suggestive as the small sample sizes limit this inference. Future studies should include broader parental age data across multiple wild populations to explore the interaction between parental age (or generation time) and *N_e_*. Larger sample sizes would also facilitate more detailed GWAS analyses and gene identification. Laboratory and common garden experiments could further explore how environmental stress influences mutation rates in externally fertilized species.

The proportion of singleton and shared mutations varies widely among taxa (Bergeron et al. 2023). We observed a high proportion of shared mutations in our data, a finding aligning with other fish DNM data (Bergeron et al. 2023; Zhang et al. 2023; Zhang et al. 2025). These recurrent mutations likely arise during the early embryonic stage before primordial germ cell specification (Jónsson et al. 2018) and are thought to be independent of parental sex bias, contributing to low paternal bias in fish mutation rates (Bergeron et al. 2023). Like the previous fish studies, we also observed no sex bias mutation rates, likely because the number of cell divisions in the two sexes in fish is thought to be more similar than in mammals and birds (Cao et al. 2021). Collectively, these observations highlight the crucial role of cell division number influencing endogenous DNA damage rates and ultimately the mutation rate in fish. The impact of replication error is especially pronounced given the short lifespan of sticklebacks, which limit the time for effective DNA repair, compounded by the higher energetic investment in reproduction.

High-quality reference genomes are essential resources for gaining an unbiased picture of genetic variation in any species. With advances in sequencing technology, the quality of *P. pungitius* genome assemblies has notably improved from version 6 (Varadharajan et al. 2019) to the telomere-to-telomere assembly presented in this paper. Lack of recombination in sex determination regions (SDR) of species with strong heterochiasmy prohibits assembly of phased sex chromosomes with the aid of linkage maps solely. Linkage map based assemblies are also error-prone when resolving complex regions enriched with structural variants and segmental duplications. These challenges were effectively solved by integrating ONT ultralong, HiFi and Hi-C data under the telomere-to-telomere (T2T) assembly framework (Li and Durbin 2024), helping us to assemble intermittent repetitive and non-repetitive regions of the chrY SDR region. Importantly, the TVA-based *NSPV9_T2T* assembly is substantially larger and more complete than earlier reference genomes, adding sequences that had not previously been assembled, including repeat rich regions on non-sex chromosomes, especially around centromeres, as well as on sex chromosomes. The absence of these regions in earlier assemblies has likely constrained research on large-scale genomic changes. As long-read sequencing technologies continue to improve, this assembly will serve as an essential resource for studying structural variation across populations.

Apart from providing the improved estimates of DNM rates in this study, the new *NSPV9_T2T* assembly provides a robust foundation for future research. Compared to calls made from an earlier assembly, fewer mutations were called from the *NSPV9_T2T* assembly with lower false discovery and false negative rates. Twenty-two mutations were detected within the previously unassembled regions (PURs), both in repetitive regions of autosomes and sex chromosomes, which are inaccessible without a gapless genome assembly. While Sanger sequencing complex genomic regions for direct verification of DNM is challenging, we observed that NSPV9_T2T reduced the false positive rate in other genomic regions improving mutation rate estimates. This may particularly improve the estimate of the localised mutation rates as the DNMs are more accurately mapped to their true genomic positions rather than to the unassembled contigs as in the previous versions. Nevertheless, as shown by validation of DNMs with Sanger sequencing, relatively high false positive rates remain a point of concern in bioinformatic DNM detection approaches even when based on best practices protocols (Wang et al. 2026b).

In conclusion, by providing the first intraspecific multipopulation test of the drift-barrier hypothesis, the results support not only its predictions but also its premises regarding the efficiency of selection in populations differing in their effective population sizes. Furthermore, the results contribute empirical evidence to the debate revolving around the relative contributions of replication errors versus DNA repair efficiency as sources of germline mutations. The lack of sex bias in mutation rates in species with similar germ cell division numbers suggest a role for replication errors (endogenous DNA damage) contributing to mutation rates. Nevertheless, the observed unique mutation spectra suggest a role also for DNA repair efficiency in contributing to mutation rates. However, we suspect that in species with short generation times such as in sticklebacks, the importance of damage rate can surpass repair efficiency in shaping the final mutation rates.

## Material and Methods

### Sampling and cross generation

This study utilized families from eight distinct populations of *P. pungitius* (see Table S6). Four populations were sampled from isolated freshwater ponds in Finland and Sweden, known to have small *N_e_*. The remaining four originated from outbred marine populations in the Baltic Sea, which have larger *N_e_* (Feng et al. 2025; see Table S6). Parental fish were collected using beach-seine nets or minnow traps during May and June 2018, then transported alive to the aquaculture facilities at the University of Helsinki. Once there, individuals from different populations were kept separately in 1 m³ plastic tanks supplied with flow-through freshwater and fed *ad libitum* twice daily with frozen chironomid larvae until they were used for artificial crosses.

Within each population, five full- or half-sib families were established through artificial fertilization, achieved by randomly pairing wild-caught individuals (Fraimout et al. 2022). As described in Zhang et al. (2023), most of these crosses involved two generations, two parents and F1 offspring, while some included a third generation (F2). Altogether, the dataset comprised 80 parental fish and 364 parent-offspring trios (see Table S6).

Standard protocols for *in vitro* fertilization and egg husbandry were followed (Barber and Arnott 2000). Eggs were obtained by gently squeezing gravid females’ abdomens over a petri dish, then fertilized by mixing with minced testes from anesthetized males (using tricaine methanesulfonate, MS-222). The fertilised eggs were kept in petri dishes with freshwater, with water changes twice daily until hatching. During this period, eggs were monitored daily for fungal infections, and any dead eggs were removed. After yolk sac absorption, fry were transferred to larger plastic boxes (approximately 11 × 10 × 10 cm) and fed live brine shrimp nauplii ad libitum. About a week later, all juveniles were moved to the Allentown Zebrafish Rack Systems (Aquaneering Inc., San Diego, CA, USA) and reared under controlled conditions with constant temperature at 15°C and a 12h:12h light-dark cycle, until they reached approximately 10 months of age (mean age: 316.4 days; sd: 23.8 days). Following the rearing period, all fish were over-anesthetized, and their fin clips preserved in 95% ethanol for subsequent DNA extraction.

### DNA extraction and sequencing

Genomic DNA was extracted from ethanol-preserved fin clips of both offspring and parents using the salting-out method described by Sunnucks and Hales (1996). The quantity and purity of DNA samples were assessed via NanoDrop spectrophotometry and Qubit™ 4.0 fluorometry (Invitrogen). DNA was available from four to five families per population (see Table S6). In addition to data from the TVA and POR populations reported in Zhang et al. (2023), all other DNA samples were sent to the Beijing Genomics Institute (BGI, Hong Kong). There, PCR-free libraries were prepared using their proprietary DNBseq platform for whole-genome resequencing, targeting an average coverage of 35×.

For telomere-to-telomere genome assembly (see Supplementary methods), a male nine-spined stickleback from the Tvärminne (TVA) population was caught in spring 2024, sacrificed and kept stored at −80°C until transported to Hong Kong in dry ice. Genomic DNA was extracted from muscle using the CTAB method in the sequencing company (Haorui Genomics, Xi’an). DNA sequencing data were generated by different platforms.The ultra-long ONT (Oxford Nanopore Technology; N50 > 100kb) reads were output from R9.4 flow cell. HiFi SMRTbell libraries were prepared using SMRTbell Express Template Prep Kit 2.0 (Pacbio, CA, USA). Then the purified products were sequenced by Revio system (Pacbio, USA) sequencer. To obtain haplotype-resolved assembly, 389× Hi-C (High-throughput Chromosome Conformation Capture) sequencing data was generated on the Illumina platform with PE150. For a high-quality genome annotation, we extracted RNA from muscle of the same individual and sequenced it on the MGISEQ-2000 platform. The species identity of this individual was confirmed by blasting the mitochondrial genome (16.58 kb in length), which was assembled with MitoHiFi (Uliano-Silva et al. 2023), against GenBank (accession OX438562.1; Figure S1).

### SNP calling, genotyping, and pedigree verification

Paired-end short-read data were aligned to both the newly assembled *NSPV9_T2T* and the Version 7 assembly (Kivikoski et al. 2021) of the nine-spined stickleback. For Version 7, due to the incomplete phasing in sex chromosome, we excluded the sex determination region (LG12: 1–16.9 Mbp) and unscaffolded contigs. Alignment files (in BAM format) were sorted with SAMtools (v1.18; Li et al. 2009) with PCR duplicates marked using Picard MarkDuplicates (v2.18; http://picard.sourceforge.net). Variants were called with GATK HaplotypeCaller (v4.3.0.0; Poplin et al. 2018) in the ERC mode, followed by base quality score recalibration (BQSR) and hard filtering according to GATK best practices (Depristo et al. 2011). The resulting gVCF files were jointly genotyped across each population with GATK’s CombineGVCFs and GenotypeGVCFs modules. Variants located within the sex determination region (SDR) were called separately, with ploidy set to 2 for females and 1 for males.

Parentage was further validated by estimating pairwise identity-by-descent (IBD) using PLINK (v1.90; Chang et al. 2015). The expected Z_0_:Z_1_:Z_2_ ratio for parent-offspring pairs is approximately 0.25:0.5:0.25. Any pedigrees showing deviations from this expectation were identified, and unfortunately, two such pedigrees from the KRK and RYT populations were excluded due to suspected false parentage, reducing the sample size for these two populations (final n_parent_ = 8, Table S6).

### Detection of candidate de novo mutations

Within parent-offspring trios, *de novo* mutations (DNM) were those deviating from Mendelian expectations. Namely, a DNM candidate was identified when the offspring possessed an allele absent from both parents. Because mutating from alternative allele to reference allele (1>0) was extremely uncommon, we limited our candidate selection to variants where both parents were homozygous for the reference (0/0) and the offspring was heterozygous (0/1). The filtering process followed the stringent criteria recommended by the GATK best practices (Poplin et al. 2018), supplemented by individual filters outlined in the “*Mutationathon*” guidelines (Bergeron et al. 2022). These filter details are provided in Figure S12 and in Zhang et al. (2023; 2025).

To reduce the likelihood of false positives, each candidate site was curated visually using IGVtools (Thorvaldsdóttir et al. 2013). Sites were discarded when the read support was inconsistent or ambiguous, where 1) reads of the alternative allele were carried by the parents or 2) poor read mapping led to incorrect heterozygous calls in the offspring. To ensure objectivity, two additional readers examined the IGV screenshots, and only sites validated by all reviewers were kept (See example in Figure S13A).

### DNM validation through Sanger sequencing

A total of 38 DNMs underwent validation through Sanger sequencing of 15 trios. Primers specific to each site were designed with Primer-BLAST (Ye et al. 2012) and ordered from Beijing Genomics Institute (BGI, Hong Kong). The polymerase chain reaction (PCR) was performed in a total volume of 50 μl for each sample, containing 50 ng of DNA template (3.5 μl), 5 μl of each primer, 25 μl of GoTaq® Green Master Mix (Promega Corporation), and 11.5 μl of deionized water. PCR was carried out using a VeritiPro Thermal Dx Cycler (Thermo Fisher Scientific Ltd, Hong Kong) on 96-well systems. The thermal cycling consisted of an initial denaturation step at 95°C for 5 minutes, followed by 30 cycles of denaturation at 95°C for 30 seconds, annealing at 60°C for 30 seconds, and extension at 72°C for 45 seconds. A final extension step was performed at 72°C for 5 minutes. The final PCR products were Sanger sequenced with paired-end reads performed by BGI (Hong Kong), and the resulting sequence traces were analyzed using Geneious(v2025.0.3; Kearse et al. 2012). A confirmed true DNM was characterized where parents were both homozygous for the reference allele showing one peak of a pure allele, while the offspring’s reads showed heterozygosity with two peaks of different alleles (see example in Figure S13B).

### Estimating de novo mutation rates

The mutation rate per generation for each individual was calculated using formula (1).

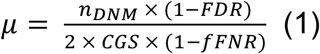

When considering the pedigree-level DNM rate (*µ*_ped_), a zero-inflated method was employed to estimate DNM rate across offspring (formula 2).

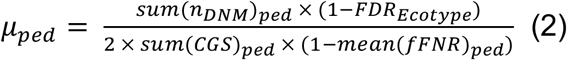

The callable genome size (CGS) was determined by counting sites within the final filtered BAM files that passed depth criteria (0.5DP_trio_ < DP_offspring_ < 2DP_trio_) and where both parents were homozygous. The false discovery rate (FDR) was derived from the proportion of candidate DNMs eliminated by PCR verification as described above, separated by two ecotypes (formula 3).

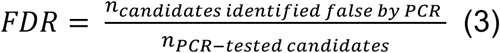

The filter false negative rate (fFNR) was inferred from the percentage of true heterozygotes in the offspring, given their parental genotypes being 0/0 and 1/1 and offspring initially identified as heterozygous (0/1), but removed due to allelic balance filtering (AB < 0.3 or AB > 0.7, formula 4).

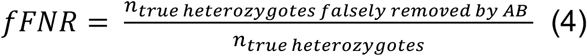

To explore parental origin bias, all DNMs were phased to identify their parent of origin using the read-backed phasing tool POOHA (https://github.com/besenbacher/POOHA). The proportion of paternal-origin mutations was calculated per pedigree to estimate the paternal-to-maternal mutation ratio (α, formula 5).

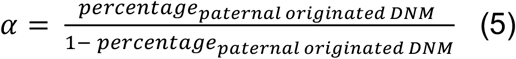

A similar formula to (2) was applied for DNM rates of the phased mutations but corrected by the pedigree-specific proportion of phasing (formula 6).

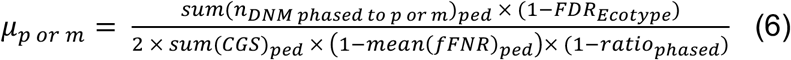

### Analysis of mutation spectrum

The mutation spectra for the candidate DNMs were analysed based on the allele changes, and they were sequentially categorised into transversions (Tv: T>G, T>A, C>A, and C>G) and transitions (Ts: T>C and C>T), or according to their pairing types into strong-to-weak (S>W: C>A, C>T), weak-to-strong (W>S: T>C, T>G), strong-to-strong (S>S: C>G), and weak-to-weak (W>W: T>A) mutations. The methylated C>T sites, inferred from the HiFi methylation data, were analysed specifically due to its high potential in deamination and mutation (Coulondre et al. 1978). The candidate DNMs were also annotated with snpEff (v5.1f; Cingolani et al. 2012) to infer the potential mutation consequences and deleterious levels of the DNMs. A specific test was performed for the DNMs that were recurrent among offspring. To ensure an equal detection probability, only the pedigree with 10 offspring were used when comparing the ratio of shared mutations.

### Age estimation

Fish age is commonly estimated by counting annual growth zones in otoliths, calcium carbonate (aragonite) structures found in the balance and hearing system in the inner ear of teleost (bony) fishes (Pannella 1971; Popper et al. 2005; Yurtseva et al. 2019). To extract the otoliths, the fish’s head was dissected by opening the cranial cavity from the top midline, allowing access to the otoliths (sagittae) located underneath the brain on the lateral sides. Cleaned and dried otoliths were placed in Eppendorf tubes and sent to collaborators at the Department of Aquatic Resources at the Swedish University of Agricultural Sciences for age determination. The otoliths were mounted on glass slides using thermosetting plastic resin (Crystalbond) and gently polished by hand in the sagittal plane with 3 µm abrasive lapping film to expose the core. The prepared sections were then stained with toluidine blue solution (0.1 g toluidine blue powder, 0.5 g NaCl, 50 ml distilled water and 0.25 ml acetic acid 100%) for two minutes, rinsed with water, and air dried prior to examination under a microscope using a combination of reflected and transmitted light. Blue-stained translucent zones were identified and counted as annuli. The ages of all lab-reared F1 parental fish were precisely known from laboratory observations.

### Estimate of effective population sizes (N_e_)

To perform a comparable analysis to Bergeron et al.(2023), we estimated *N_e_* using the Pairwise Sequentially Markovian Coalescence (PSMC) method (Li and Durbin 2024). We randomly chose one wild-caught male sample from each population, and the raw fastq sequences were converted into fasta format with fq2psmcfa, setting the minimum depth to one-third of the sample’s average coverage and the maximum depth to twice. The parameters were configured as follows: –N25 for the maximum number of iterations, –t15 as the upper bound for the time to the most recent common ancestor, – r5 for the initial θ/ρ value, and the atomic interval pattern –p of ‘4 + 25 × 2 + 4 + 6’. The generation time was set to 2 years for marine populations and 3 years for freshwater populations according to DeFaveri et al. (2014). Mutation rates estimated from this study were applied in the analysis. The harmonic mean over windows spanning from 30,000 to 1,000,000 years was calculated and used in the final estimates.

Additionally, we estimated the contemporary *N_e_* using CurrentNe2 (Santiago et al. 2025), which relies on linkage disequilibrium (LD) patterns between SNPs without requiring their specific genomic locations or a genetic map. This method also accounts for the subpopulation structure by allowing migration (-x).The VCF files were randomly thinned to retain only one SNP per 10bp using VCFtools (--thin 10; v 0.1.16; Danecek et al. 2011) resulting in VCF files below 2Mb SNPs, and the recombination rates applied in this analysis were from a manuscript in preparation.

### Estimate of selection strength

To test if the small freshwater populations have experienced reduced selection as compared to larger marine populations, the toolkit “degenotate” embedded with snpArcher(Mirchandani et al. 2024) were used to perform MacDonald–Kreitman (MK) test for each gene (Mcdonald and Kreitman 1991). The outgroup used in this analysis was from RUS-LEV (66.3°N, 33.4°E) which was known to be an ancestral population of the studied populations belonging to the Eastern European lineage of this species (Feng et al. 2024). The overall neutrality index (*Ni*) for population was calculated following the formula (7) where P_n_ and P_s_ were the number of nonsynonymous and synonymous polymorphisms, whereas D_n_ and D_s_ were those substitutions fixed in each population.

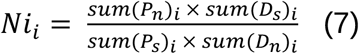

An *Ni* value above one indicates purifying selection, while a value less than one suggests positive selection.

To further evaluate selection efficacy, we utilised the slope of the linear relationship between nucleotide diversity and the recombination rate. The ability of recombination to maintain nucleotide diversity is theoretically constrained in populations where genetic drift is dominant and selection is inefficient (Wang et al. 2026a). Under such conditions, the ‘diversity-recombination’ slope flattens, as stochastic processes overpower the signature of linked selection.

### Statistical analyses

Phylogenetic generalized linear mixed models (PGLMM) were constructed using the R package *MCMCglmm* (v 2.36; Hadfield 2010) to investigate the relationship between per-generation DNM rates and *N_e_* estimates from both PSMC and CurrentNe2. Additionally, the models assessed the association between phased DNM rates and parental age, incorporating genetic relatedness among samples as a random effect. The phylogeny was inferred with IQtree2 (v 2.3.6; Minh et al. 2020) by randomly selecting one offspring from each pedigree, followed by standard model selection, tree inference, and 3,000 bootstrap replicates using the Optimize UFBoot method with NNI on the bootstrap alignment. The resulting phylogenetic tree is provided in the supplementary material.

The first model analysed DNM counts using a quasipoisson distribution (Family = “poisson”), with an offset based on the scaled number of callable genome sites (CGS), corrected for false discovery rate (FDR) and false negative rate (FNR). The model formula was:

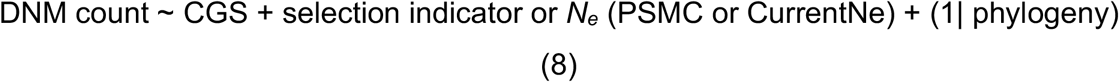

Similarly, a bivariate model was employed to test the joint effect of parental age to the per generation or per year DNM rates, similarly to the above model settings:

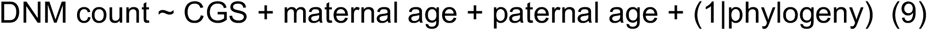

Another PGLMM model was applied to test the effect of parental age on phased DNM rates using Gaussian distribution, separately for maternal and paternal sides, incorporating if they were recurrent mutations among offspring:

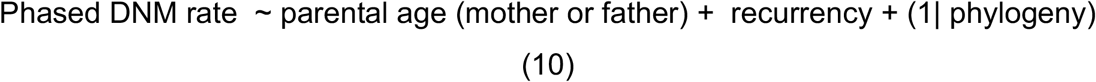

All PGLMM models used a chain length of 1,000,000 iterations, with a burn-in of 200,000 and a thinning of 100, which resulted in 8,000 posterior samples. Convergence was assessed by examining plots of fixed and random effects.

Furthermore, by integrating data from other vertebrates (Bergeron et al. 2023), we fitted generalized linear mixed models (GLMM) to examine the relationships between DNM rates, either per year or per generation, and variables such as *N_e_* or generation time (GT). These models were fitted using the R package *glmmTMB* (v 1.1.11; Brooks et al. 2017) with a Gaussian distribution, incorporating pedigree as a random effect. DNM rate and *N_e_* were log scale adjusted.

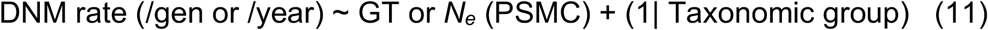

A GLMM model was also fitted in *glmmTMB* to assess the correlation between *Ni* or π/r slope and the per-generation DNM rates. The model employed a Poisson distribution weighted by the normalised size of callable genome:

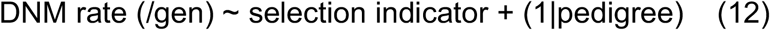

A similar approach were applied to test the joint impacts from the parental ages in a bivariate model:

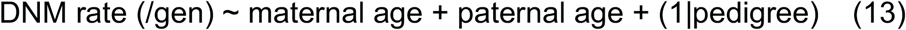

The variance explained by these models was calculated using R package *MuMIn* (v1.48.11; Kamil 2025). All models were checked by plotting residuals against fitted values, and dispersion estimates were examined to avoid overdispersion.

### Genome wide association analyses

To evaluate the association between parental genetic variants and the pedigree-level DNM rate (*µ*_ped_) or the phased DNM rate, genome-wide association studies (GWAS) were performed. The analysis utilised three different statistical models: the General Linear Model (GLM), the Mixed Linear Model (MLM), and the Fixed and Random Model Circulating Probability Unification (FarmCPU) in rMVP (Yin et al. 2021). To reduce the confounding effects from genetic relatedness, the first five principal components derived from the SNP matrix were included as covariates in the analyses. The significance of associations was determined using a Bonferroni correction, dividing by the total number of SNPs tested, and the adjusted threshold was set to 0.05. To investigate the functions of the proteins where the associated SNPs were situated, their sequences were blasted against the UniProt database (https://www.uniprot.org/). The same sequences were also utilised in pathway enrichment analyses in clusterProfiler (Yu et al. 2012).

## Acknowledgements

We appreciate Dr. Antoine Fraimout for collecting the samples and developing the pedigrees for this project. We thank the support from Miinastiina Issakainen and Kirsi Kähkönen for their assistance with DNA extractions. We are also thankful to Nick Ho Ming Lin, Zahra, Hyein Kil, and Mei Ying Ng for their help in PCR experiments. We are grateful to Dr. Ulrika Candolin in helping obtain the specimen used for T2T genome assembly. Thanks also to Dr. Xueyun Feng, Dr. Jilong Ma and Joanna Maesel for their valuable advice. Our research was supported by Seed Fund for Basic Research (#2309100132) and General Research Fund (#17104824 and #17126824) from Research Grants Council (Hong Kong), and Academy of Finland (#218343 to JM). We acknowledge the Finnish IT Centre for Scientific Computing (CSC) for access to computational resources.

## Ethic statement

The fish breeding was conducted under a permit from the Animal Experiment Board in Finland (permit reference ESAVI/4979/2018). The fish of the parental generation were collected under national fishing licences.

## Author contributions

Conceived and designed the study: JM, CZ, MHS

Reference genome assembly: HW, DW

Mutation identification and data analyses: CZ, KR, LL

Wrote the paper: CZ, JM, DW, HW, KR, MHS

## Data accessibility statement

The NSPV9_T2T genome assembly of *P. pungitius* has been deposited in the National Center for Biotechnology Information (NCBI; https://www.ncbi.nlm.nih.gov/) at PRJNA1331522. The raw whole-genome sequencing data will be provided in NCBI under accession code PRJNA1334546 at the time of acceptance. The phylogenetic tree applied in the phylogenetic generalized linear mixed models is available in Supplementary data.

