## Supplementary for "Intra-specific evidence to support the drift-barrier hypothesis of mutation rate evolution"

**Supplementary information**

Table of contents

**Supplementary methods**

**Supplementary Tables:**

Supplementary Table 1. Genome sequencing platforms and depth for *NSPV9_T2T*.

Supplementary Table 2. Genome completeness of *NSPV9_T2T* by BUSCO assessment.

Supplementary Table 3. Number of annotated features using Eukaryotic Genome Annotation Pipeline.

Supplementary Table 4. Completeness of annotated proteins in *NSPV9_T2T* by BUSCO assessment.

Supplementary Table 5. Number of SNPs passed each DNM filter and the mean values for the rate components.

Supplementary Table 6. Sampling information and rate of germline *de novo* mutations (DNM) in the eight populations.

Supplementary Table 7. Generalized mixed linear model results for de novo mutation rates.

**Supplementary Figures:**

Supplementary Figure 1. Alignment plot for the mitochondrial genomes of *NSPV9_T2T* and OX438562.1 (Genebank ID).

Supplementary Figure 2. Hi-C interaction heatmap for haplotype 1 of *Ppn-T2T*.

Supplementary Figure 3. Hi-C interaction heatmap for chrY, haplotype 2.

Supplementary Figure 4. Sequencing collinearity between version 7, *NSPV9_T2T* and version 8 assembly in *Pungitius pungitius*.

Supplementary Figure 5. Ecotype-comparison of the per-generation and per-year germline *de novo* mutation rates at the individual level.

Supplementary Figure 6. Correlation between per-generation rates of DNMs and the coalescent and contemporary effective population sizes (N_e_).

Supplementary Figure 7. Correlation between the per-year DNM rates and harmonic mean of effective population sizes (*N_e_*) estimated from PSMC across vertebrates.

Supplementary Figure 8. Population selection strength and its correlation with per-generation DNM rates.

Supplementary Figure 9. Ecotype-comparison of the pedigree-level Ts/Tv and S>W/W>S ratio

Supplementary Figure 10. Results from the genome-wide association study (GWAS) and function enrichment analyses.

Supplementary Figure 11. A potential structure variant found located on a long non-coding RNA (*lncrna*) in the hypermutable pedigree fin_ryt22.

Supplementary Figure 12. Illustration of the *de novo* mutation occurring at different chromosomes and the filtering setting based on the ploidy.

Supplementary Figure 13. DNM verification examples

**Supplementary Methods**

Genome assembly and Y chromosome scaffolding

We performed a hybrid assembly using Hifiasm (v0.20.0‑r639; Cheng et al. 2024) and Verkko (v2.2.1; Rautiainen et al. 2023), integrating PacBio HiFi, ultra‑long ONT, and Hi‑C sequencing data. To improve the continuity of the telomeric regions, telomeric motifs (--telo‑m CCCTAA) were provided to both tools, and the Hifiasm contigs were further assembled based on N50. To make sure all contigs were from the reference individual nuclear genome, the mitochondrial genome was assembled and annotated with MitoHiFi (v3.2.2; Uliano-Silva et al. 2023), after which redundant contigs were removed. One contaminated nuclear genome contig, likely derived from *Gyrodactylus salaris*, was identified via BLASTn (BLAST+, v2.16.0; Camacho et al. 2009) against the NCBI nucleotide collection (nt; accessed on 2024-10-29) database and excluded. Hi‑C data were mapped and filtered to the draft genome following the Arima‑HiC pipeline (https://github.com/ArimaGenomics/mapping_pipeline). The contig assembly was scaffolded to the chromosomal level with YaHS (v1.2.2; Zhou et al. 2023) based on the Hi-C signals.

Due to highly repetitive elements on the large chrY, the contigs were still fragmented. Thus the orientation and the order of these Y-contigs were further manually corrected with JuiceBox (v1.11.08; Durand et al. 2016). Five high-quality contigs in sex determination region (SDR) were generated from Hifiasm, and these were reanchored according to the Verkko results, which output one contiguous Y-contig.

Gap filling and genome polishing

Assembly gaps, especially those involving segmental duplications (SDs), were filled with ultra-long ONT sequences. assembled with NextDenovo (v2.5.2; Hu et al. 2024a) and Flye (2.9.5-b1801; Kolmogorov et al. 2019) as well as the combined data (HiFi + ONT + Hi-C) with Verkko. They were then polished using HiFi reads in Hypo (v1.0.3; Ritu et al. 2019). The RagTag patch model was applied to close all gaps in both haplotype assemblies, except for one gap in chrY which was addressed with ultra-long ONT reads using TGS-GapCloser2 (Xu et al. 2020). Genome polishing was performed in three rounds. In the first round, variants were called respectively with HiFi reads in DeepVariant (Poplin et al. 2018), ONT reads in Clair3 (v1.0.10; Zheng et al. 2025), and the haplotype-resolved assembly generated by Verkko using PAV (v2.4.6; Ebert et al. 2021), where the overlapped alternative homozygotes (1/1) were considered as potential errors. Secondly, errors identified in Merqury (v1.3; Rhie et al. 2020) were then polished using the “Polisher” model in Flye. Finally, a last round of polishing was performed with HiFi alignment in NextPolish2 (v0.2.1; Hu et al. 2024b).

Annotation of the repeats and genes

To comprehensively characterise the repetitive elements within the genome, we adopted a combined annotation strategy of homology-based searching and *de novo* prediction. RepeatMasker (v4.1.5; Tarailo-Graovac & Chen) was firstly used to search homologs in the RepBase (v21.12; Jurka et al. 2005) and Dfam (v3.7; Storer et al. 2021) databases for known repeat families, and then used to search in the *de novo* library constructed by RepeatModeler2 (v2.0.4; Flynn et al. 2020). Regarding the gene annotation, the NCBI Eukaryotic Genome Annotation Pipeline (EGAPx, v0.4.0-alpha; https://github.com/ncbi/egapx) was utilised to annotate genes based on transcriptome and protein evidence. We integrated mixed RNA-seq data from the reference individual, four new sequenced samples (Wang et al. *in preparation*), 48 public nine-spined stickleback samples (Wang et al. 2020), and Iso-seq long-read transcript data (study accession: PRJEB80638), to enhance the accuracy and completeness of the annotation.

Genome Quality and Completeness Estimate

We evaluated the assembly quality from two aspects: the completeness and the base accuracy (quality value, QV). The BUSCO (v5.8.2; Simão et al. 2015) was used to assess the completeness of core single-copy orthologous genes in Actinactinopterygii lineage. Merqury was applied for a k-mer-based consensus assessment of the assembly's base accuracy. This method calculated the QV score by comparing k-mers from the assembly with HiFi reads, accounting for k-mers that exist only in the assembly as potential errors, while those presented in both the assembly and the raw data as correct validation.

Previously Unassembled Regions (PURs) Identification

To identify sequence regions potentially missing in the previous assembly versions but present in *NSPV9_T2T*, we performed a PUR analysis. We first aligned the Version 7 assembly to *NSPV9_T2T* using Winnowmap2 (v2.03; Jain et al. 2022), and then we performed format conversion and quality filtering to obtain high-quality alignment fragments. Subsequently, these fragments were merged based on their coordinates on the reference genome to create a map of covered regions. Finally, we used the ‘complement’ function of bedtools (v2.31.1; Quinlan & Hall 2010) to locate the uncovered segments on the reference genome, defining them as PURs.

**Supplementary Table 1.** Genome sequencing platforms and depth for *NSPV9_T2T*.

| **Reads** | **Depth** | **Platforms** |
| --- | --- | --- |
| Oxford Nanopore (ONT) long reads | 84.6× | Oxford Nanopore PromethION |
| PacBio HiFi long reads | 173.8x | PacBio Revio system |
| Hi-C reads (DpnII restriction enzyme) | 389.0× | MGISEQ-2000 |

**Supplementary Table 2.** Genome completeness of *NSPV9_T2T* by BUSCO assessment.

| **Complete BUSCOs** | **Complete and single-copy BUSCOs** | **Complete and duplicated BUSCOs** | **Fragmented BUSCOs** | **Missing BUSCOs** | **Total Lineage BUSCOs** |
| --- | --- | --- | --- | --- | --- |
| 3603  (99.0%) | 3345  (91.9%) | 258  (7.1%) | 22  (0.6%) | 15  (0.4%) | 3,640  (100%) |

**Supplementary Table 3.** Number of annotated features using Eukaryotic Genome Annotation Pipeline.

| Genes | 23760 |
| --- | --- |
| genes (other) | 0 |
| genes (non-transcribed pseudo) | 232 |
| genes (transcribed pseudo) | 0 |
| genes (has variants) | 7359 |
| genes (partial) | 30 |
| genes (Ig TCR segment) | 13 |
| genes (non coding) | 552 |
| genes (protein coding) | 22963 |
| genes (major correction) | 254 |
| genes (minor correction) | 0 |
| genes (premature stop) | 39 |
| genes (has frameshifts) | 229 |
| mRNAs | 37293 |
| mRNAs (exon <= 3nt) | 10 |
| mRNAs (partial) | 30 |
| mRNAs (correction) | 254 |
| mRNAs (model) | 37293 |
| mRNAs (fully supported) | 35737 |
| mRNAs (ab initio > 5%) | 775 |
| mRNAs (has gaps) | 0 |
| mRNAs (model with correction) | 254 |
| Non-coding RNAs | 1473 |
| non-coding RNAs (exon <= 3nt) | 0 |
| non-coding RNAs (partial) | 0 |
| non-coding RNAs (correction) | 0 |
| non-coding RNAs (model) | 1473 |
| non-coding RNAs (fully supported) | 1473 |
| non-coding RNAs (ab initio > 5%) | 0 |
| non-coding RNAs (has gaps) | 0 |
| Pseudo transcripts | 232 |
| pseudo transcripts (exon <= 3nt) | 0 |
| pseudo transcripts (partial) | 0 |
| pseudo transcripts (correction) | 0 |
| pseudo transcripts (model) | 232 |
| pseudo transcripts (fully supported) | 110 |
| pseudo transcripts (ab initio > 5%) | 0 |
| pseudo transcripts (has gaps) | 0 |
| Coding sequences (CDSs) | 37293 |
| CDSs (exon <= 3nt) | 656 |
| CDSs (partial) | 30 |
| CDSs (correction) | 254 |
| CDSs (model) | 37293 |
| CDSs (fully supported) | 35737 |
| CDSs (ab initio > 5%) | 878 |
| CDSs (model with correction) | 254 |
| CDSs (major correction) | 254 |
| CDSs (minor correction) | 0 |
| CDSs (premature stop) | 39 |
| CDSs (has frameshifts) | 229 |

**Supplementary Table 4.** Completeness of annotated proteins in *NSPV9_T2T* by BUSCO assessment.

| **Complete BUSCOs** | **Complete and single-copy BUSCOs** | **Complete and duplicated BUSCOs** | **Fragmented BUSCOs** | **Missing BUSCOs** | **Total Lineage BUSCOs** |
| --- | --- | --- | --- | --- | --- |
| 3607(99.1%) | 3340 (91.8%) | 267 (7.3%) | 13 (0.4%) | 20 (0.5%) | 3640 (100%) |

**Supplementary Table 5.** Number of SNPs passed by each DNM filter and the mean values for the rate components. Column names are the population names with the number of parent-offspring trios in parentheses.

A. T2T assembly (filter stats including SDR)

| **Filter** | **TVA**  **(52)** | **POR**  **(54)** | **UME**  **(58)** | **RAA**  **(54)** | **BYN**  **(33)** | **PYO**  **(49)** | **RYT**  **(40)** | **KRK**  **(24)** |
| --- | --- | --- | --- | --- | --- | --- | --- | --- |
| Site filtering | 4328219 | 4478188 | 4459989 | 4522193 | 1882389 | 2299619 | 1925681 | 1745710 |
| Mendelian Violation +Individual filtering | 10202 | 8835 | 8116 | 7314 | 1506 | 2660 | 2712 | 542 |
| Repetitive between pedigrees | 513 | 443 | 495 | 377 | 159 | 248 | 246 | 44 |
| Manual Curation | 118 | 106 | 151 | 88 | 134 | 116 | 191 | 31 |
| **Other rate components** | |  |  |  |  |  |  |  |
| Mean CGS (autosome) | 358.04 | 346.17 | 332.71 | 345.34 | 346.64 | 344.99 | 344.19 | 342.96 |
| Mean FDR | 21.05% | 21.05% | 21.05% | 21.05% | 10.00% | 10.00% | 10.00% | 10.00% |
| Mean FNR | 5.47% | 5.73% | 7.10% | 6.11% | 5.80% | 6.86% | 5.37% | 5.39% |

B. Version 7 (filter stats excluding SDR)

| **Filter** | **TVA**  **(52)** | **POR**  **(54)** | **UME**  **(58)** | **RAA**  **(54)** | **BYN**  **(33)** | **PYO**  **(49)** | **RYT**  **(40)** | **KRK**  **(24)** |
| --- | --- | --- | --- | --- | --- | --- | --- | --- |
| Site filtering +Mendelian Violation | 506121 | 660606 | 621481 | 639810 | 178241 | 80400 | 161746 | 137890 |
| Individual filtering | 3462 | 3857 | 4233 | 3128 | 735 | 733 | 1324 | 366 |
| Repetitive between pedigrees | 931 | 995 | 1002 | 907 | 198 | 243 | 377 | 100 |
| Manual Curation | 106 | 121 | 148 | 91 | 130 | 154 | 201 | 31 |
| **Other rate components** | |  |  |  |  |  |  |  |
| Mean CGS (autosome) | 357.97 | 346.13 | 330.56 | 345.32 | 336.39 | 363.66 | 354.33 | 349.07 |
| Mean FDR | 17.65% | 17.65% | 17.65% | 17.65% | 23.08% | 23.08% | 23.08% | 23.08% |
| Mean FNR | 5.64% | 5.96% | 7.61% | 6.32% | 6.21% | 6.64% | 5.76% | 5.90% |

**Supplementary Table 6.** Information on sampling locations, estimates of effective population size applying PSMC and CurrentNe2 (detailed in Material and Methods), neutrality index, sample sizes, number of de novo mutations (DNM) observed in each offspring and rate of DNM estimated in the eight populations.

| **Population** | **Code** | **Coordinate** | **Ecotype** | ***Ne*_PSMC-hm_** | ***Ne_CurrentNe2_*** | **Neutrality Index** | **n_family_** | **n_trio_** | **Mean**  **#DNM** | **DNM rate**  **(10^-9^/gen/bp)** |
| --- | --- | --- | --- | --- | --- | --- | --- | --- | --- | --- |
| Pori | POR | 61.59°N, 21.47°E | MA | 163297 | 12972.6 | 1.59 | 5 | 54 | 1.80 | 2.51 |
| Raahe | RAA | 64.69°N, 24.46°E | MA | 211821 | 10925.9 | 1.65 | 5 | 54 | 1.56 | 2.18 |
| Tvärminne | TVA | 59.83°N, 23.0°E | MA | 152778 | 24262.0 | 1.54 | 5 | 52 | 2.06 | 2.77 |
| Umeå | UME | 63.64°N, 19.99°E | MA | 127591 | 9136.8 | 1.56 | 5 | 58 | 2.36 | 3.48 |
| Bynastjärnen | BYN | 64.45°N, 19.44°E | FW | 33140 | 403.9 | 1.43 | 5 | 33 | 3.58 | 5.70 |
| Kirkasvetinenlampi | KRK | 66.44°N, 29.14°E | FW | 32767 | 266.5 | 1.23 | 4 | 24 | 1.00 | 1.60 |
| Pyöreälampi | PYO | 66.26°N, 29.43°E | FW | 25988 | 18.4 | 1.39 | 5 | 49 | 2.18 | 3.52 |
| Rytilampi | RYT | 66.38°N, 29.32°E | FW | 38253 | 384.0 | 1.20 | 4 | 40 | 4.38 | 6.99 |

**Supplementary Table 7. Generalized mixed linear model results for de novo mutation rates.**

A. MCMCglmm models with the pedigree-level phylogeny as the random effect.

| Response variable | Random effect | Fixed effects | Slope | p-value | DIC |
| --- | --- | --- | --- | --- | --- |
| #DNM  (indv-level; Quaispoisson distribution; CGS corrected by FNR, FDR and normalised; *N_e_* were log-scaled) | Pedigree  (phylogeny)  <1% | CurrentNe2  +CS_scaled  intercept | -0.0923  0.1607  1.5416 | .0703  .0343*  .0043** | 1003.339 |
|  |  | PSMC2_hm_  +CS_scaled  intercept | -0.3986  0.2077  5.3549 | .0075**  .0075**  .0025** | 1001.702 |
|  |  | Age_average_  +CS_scaled  intercept | 0.0312  0.1201  0.7558 | .7797  .0915  .0370* | 1092.792 |
|  |  | Neutrality Index  +CS_scaled  intercept | -0.1206  0.1196  1.0111 | .8790  .1230  .4010 | 1007.527 |
| rate/gen  (fam-level; Normal distribution; corrected) | Pedigree  (Phylogeny)  8.92% | Ecotype Intercept | -0.7700  0.2084 | .00325**  .07600 | 206.324 |
| rate/gen  (fam-level; Normal distribution; corrected) | Pedigree  (Phylogeny)  21.17% | PM (PM)  Intercept | -0.8165  0.3301 | <1e-04***  .328 | 180.004 |
| rate/gen **- Maternal**  (fam-level; Normal distribution; corrected) | Pedigree  (Phylogeny)  41.18% | Age  +PM (PM)  Intercept | 0.0370  -0.6505  0.0669 | .7775  .0005***  .8922 | 88.120 |
| rate/gen **- Paternal**  (fam-level; Normal distribution; corrected) | Pedigree  (Phylogeny)  33.06% | Age  +PM (PM)  Intercept | 0.0388  -0.8744  0.2134 | .8190  <1e-04***  .6370 | 95.409 |

B. glmmTMB models with taxonomic group as the random effect

| Response variable | Random effect | Fixed effects | Slope | p-value | AIC |
| --- | --- | --- | --- | --- | --- |
| Per-gen mu  (log-scale; Normal distribution) | Taxonomic group  13.68% | PSMC2_hm_ (log-scaled)  Intercept  7.10% | -0.0918  -17.9157 | 0.0234*  <2e-16*** | 85.1 |
| Per-year mu  (log-scale; Normal distribution) | Taxonomic group  (2.99%) | PSMC2_hm_ (log-scaled)  Intercept  (34.00%) | 0.4298  -25.1350 | 5.52e-10***  <2e-16*** | 197.1 |
|  | Taxonomic group  (0.00%) | Generation time  Intercept  (47.17%) | -0.1075  -19.5697 | <2e-16***  <2e-16*** | 181.0 |

**Supplementary Figure 1.** Alignment plot for the mitochondrial genomes of *NSPV9_T2T* and OX438562.1 (Genebank ID).

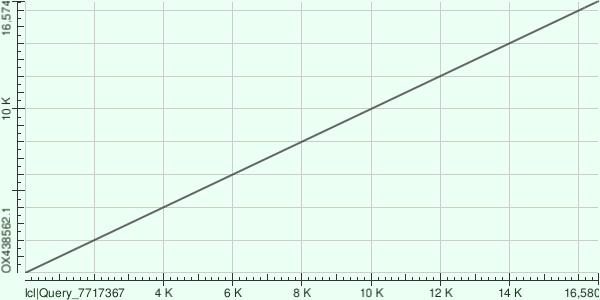

**Supplementary Figure 2.** Hi-C interaction heatmap for haplotype 1 of *NSPV9-T2T*. The clear diagonal signal confirms the continuity and accuracy of the sequence assembly.

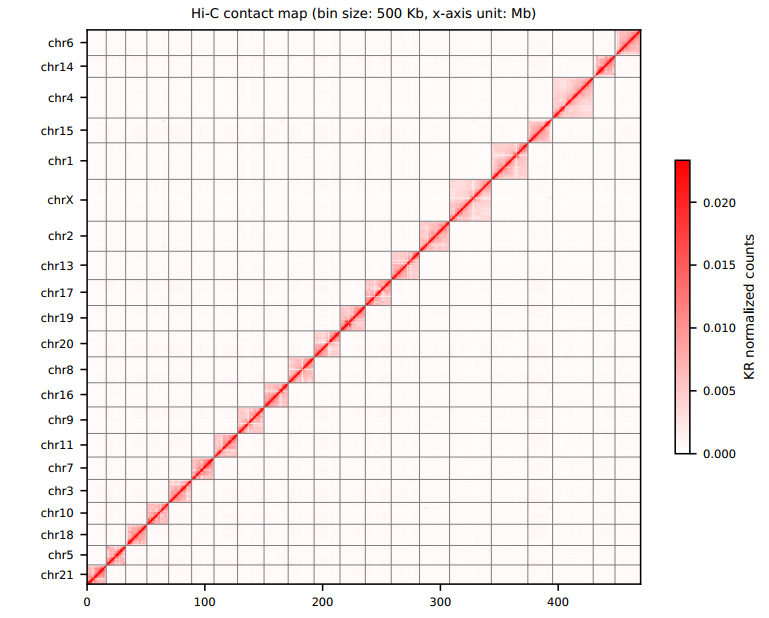

**Supplementary Figure 3.** Hi-C interaction heatmap for chrY, haplotype 2 of *NSPV9_T2T*. The strong diagonal signal confirms the continuity and accuracy of chrY, although the sex determination region remains unidentified in this short-read built plot.

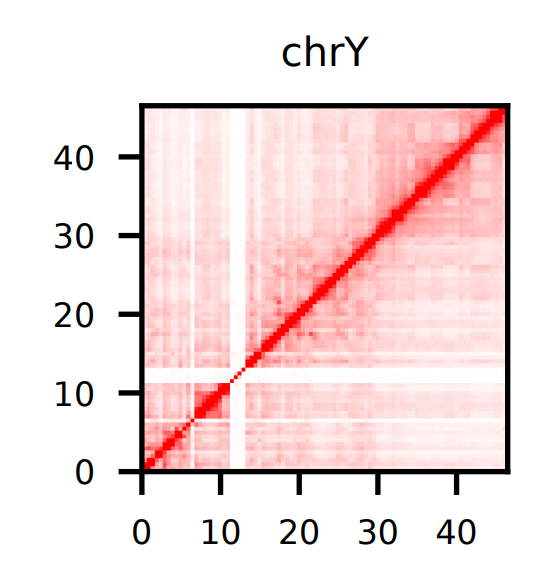

**Supplementary Figure 4.** Sequencing collinearity between version 7, *NSPV9_T2T* and version 8 assembly of *Pungitius pungitius genome*, centred by *NSPV9_T2T*.

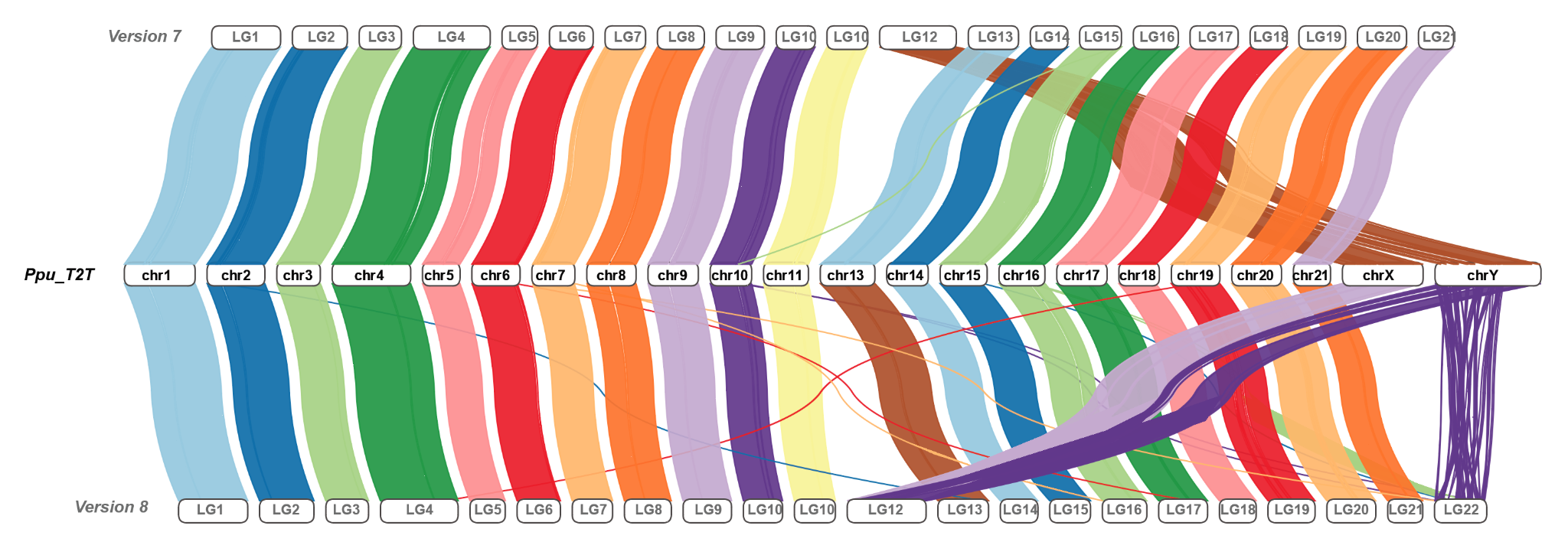

**Supplementary Figure 5.** Ecotype-comparison of the (A) per-generation and (B) per-year germline *de novo* mutation rates at the individual level. Individuals from the hypermutable pedigree, *fin_ryt22*, were colored in red and excluded from the t-tests.

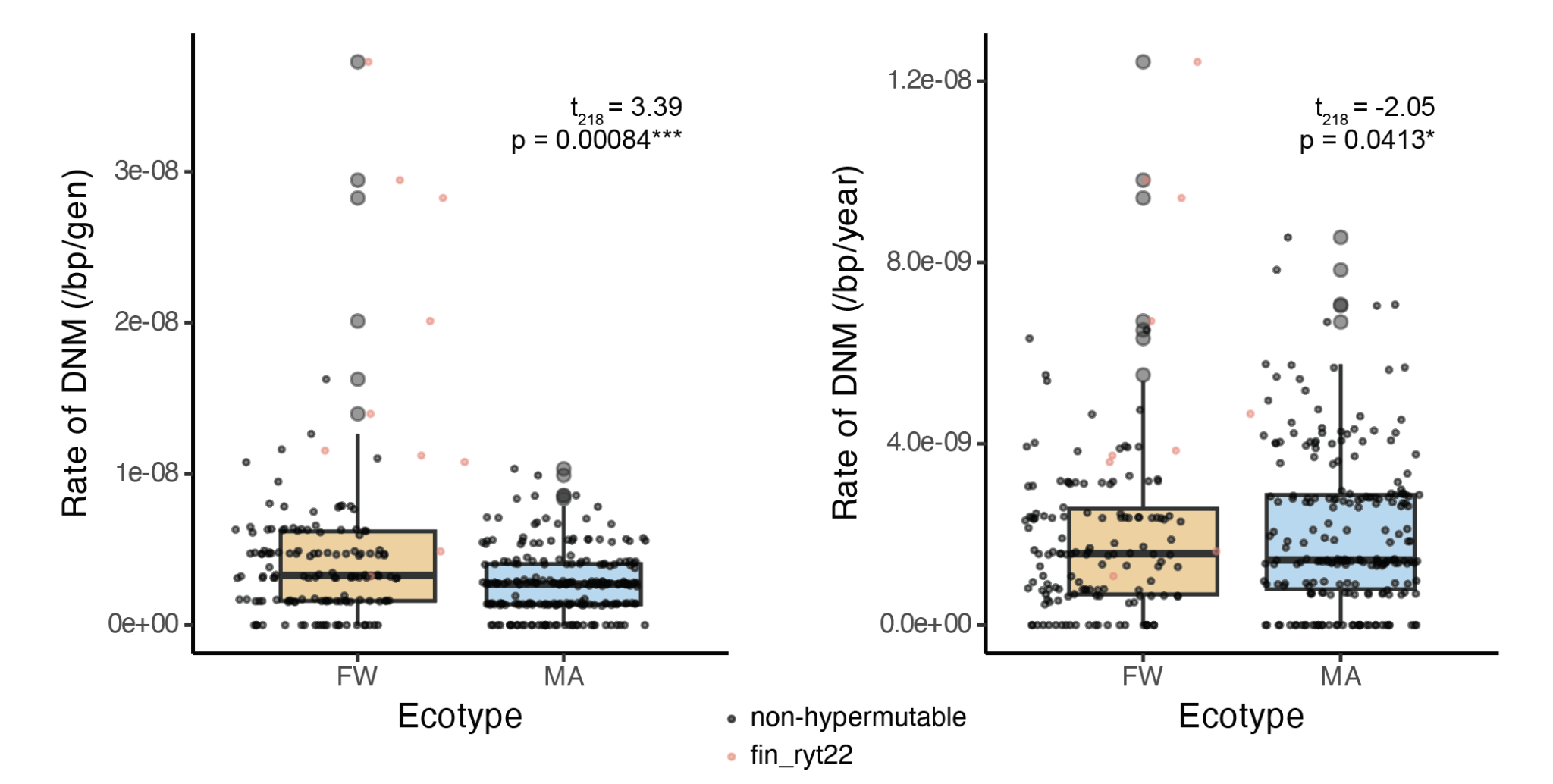

**Supplementary Figure 6.** Correlation between pedigree-level rates of DNMs (/generation/bp) and A) the harmonic mean of coalescent PSMC effective population sizes (N_e_), or contemporary N_e_ estimated from CurrentNe2. P-values were from a phylogenetic generalised linear mixed model using MCMCglmm, excluding the outlier hypermutable pedigree. Model details please see *Material and Methods*.

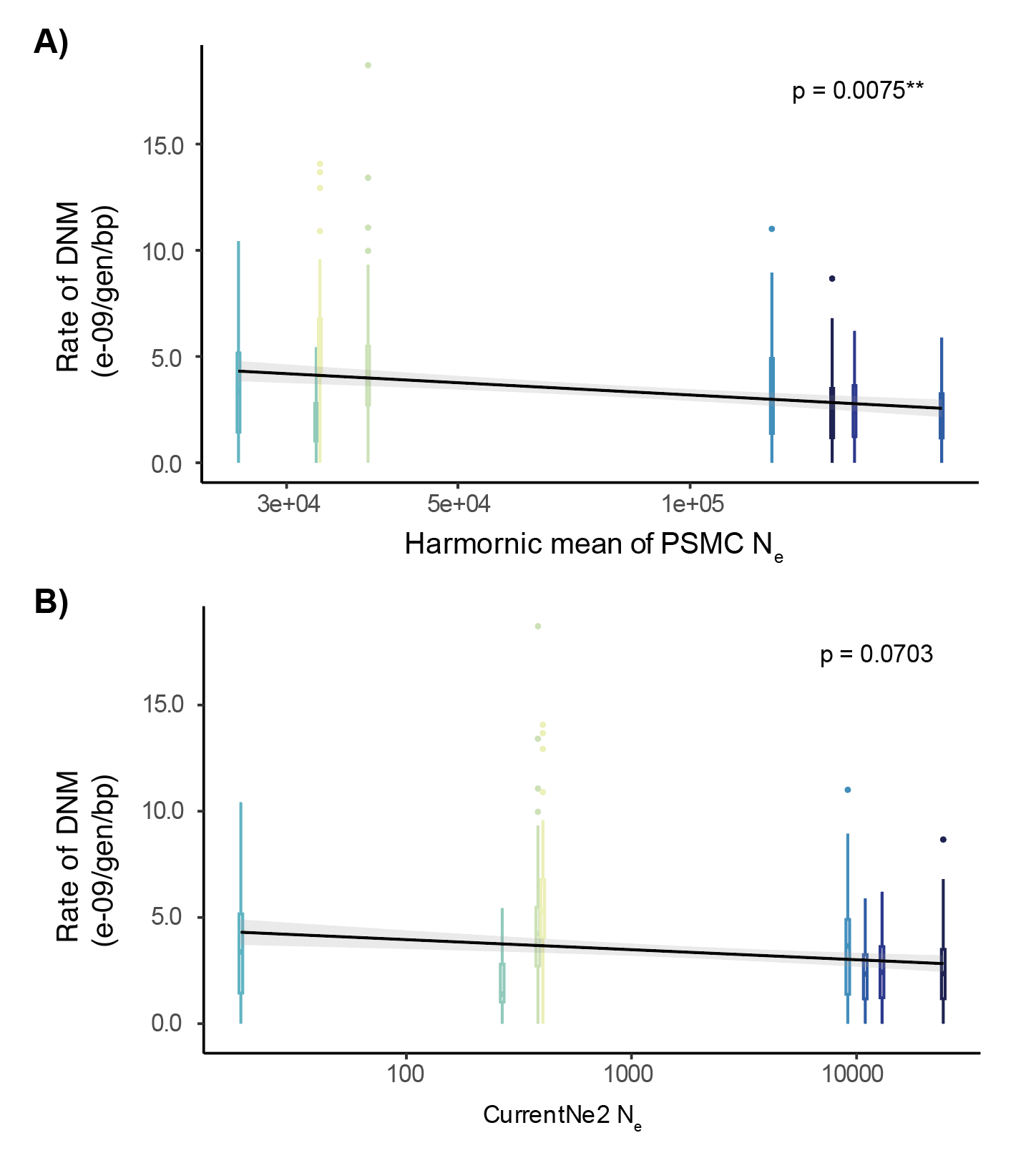

**Supplementary Figure 7.** Correlation between the per-year DNM rates and harmonic mean of effective population sizes (*N_e_*) estimated from PSMC across vertebrates.

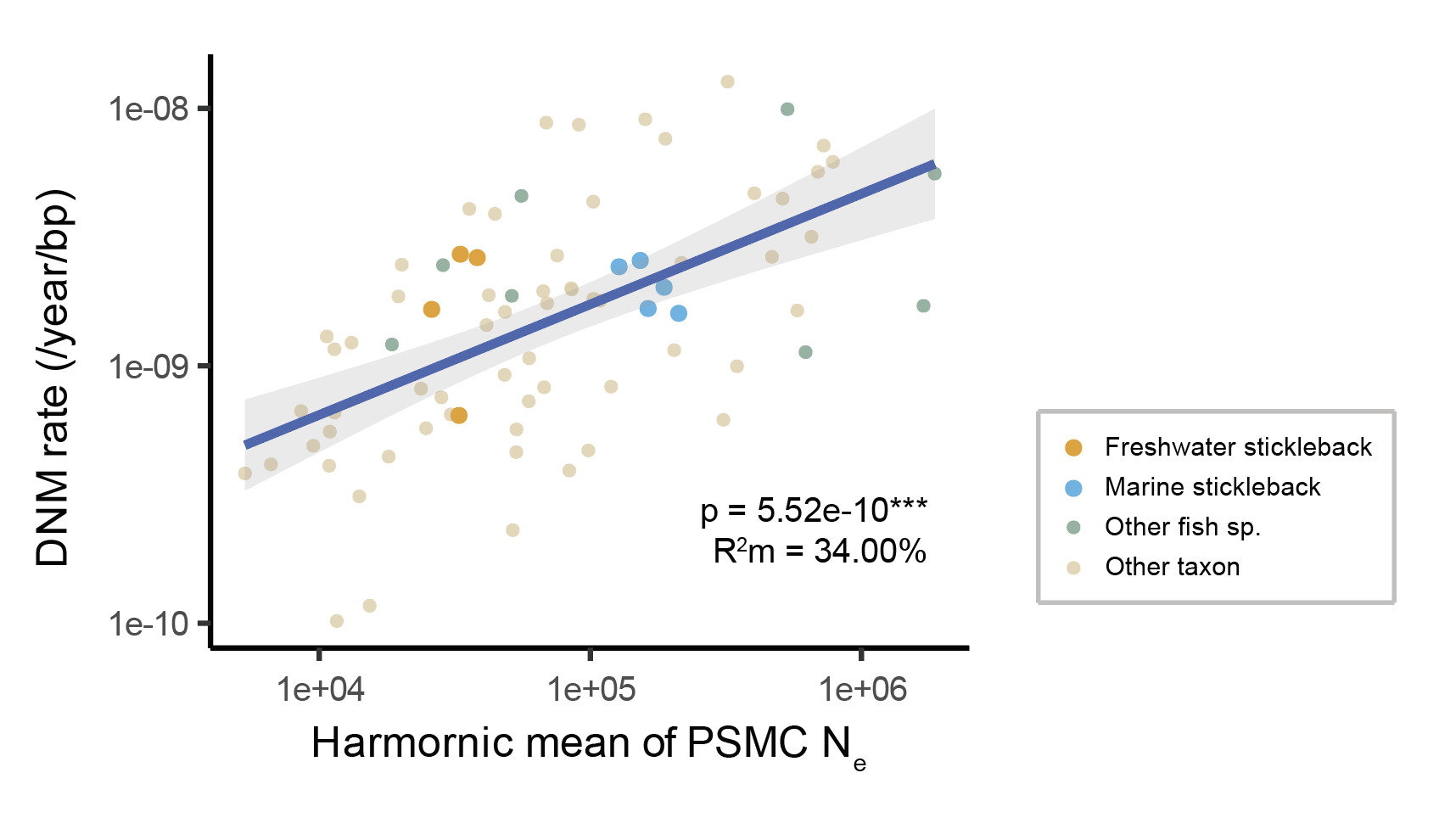

**Supplementary Figure 8.** Selection strength and its association with per-generation DNM rates in different populations and ecotypes. A) neutrality indexes (*Ni*) of the negatively-selected genes in the eight populations estimated in the McDonald–Kreitman test. B) correlation between the overall population Ni and the per-generation DNM rates.

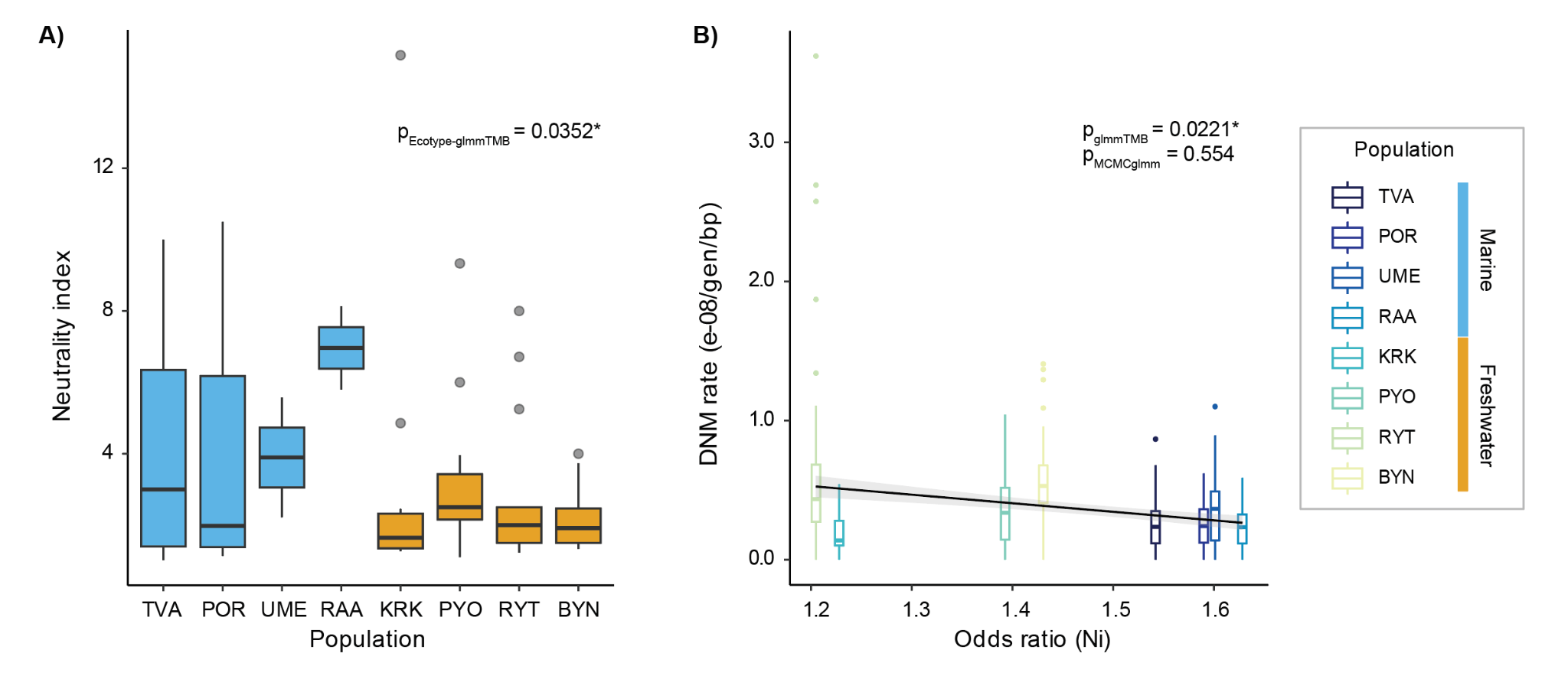

**Supplementary Figure 9.** Ecotype-comparison of the pedigree-level A) transition to transversion (Ts/Tv) and B) strong-to-weak/weak-to-strong (S>W/W>S) ratio.

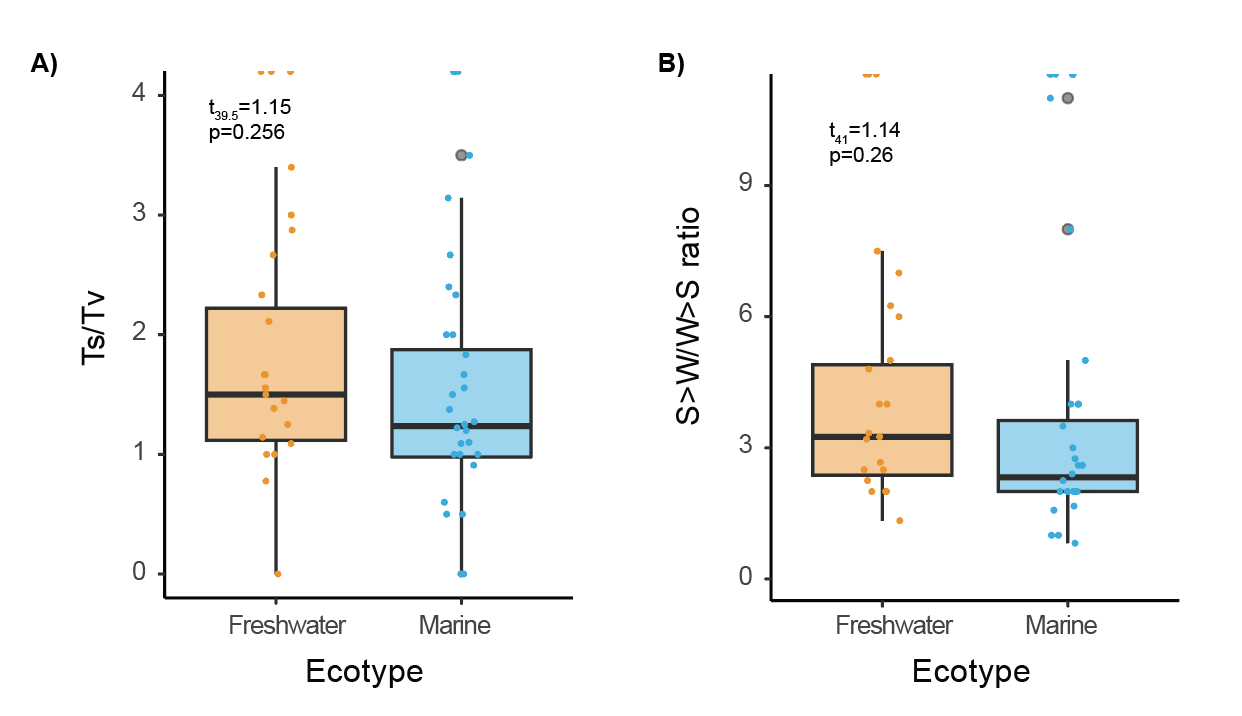

**Supplementary Figure 10.** Results of the genome-wide association study (GWAS) for A) the phased DNM rates (different between two parents), B) the pedigree-level DNM rates (the same for the two parents). C) the result from pathway enrichment analysis with the MLM model, combining the results from two traits.

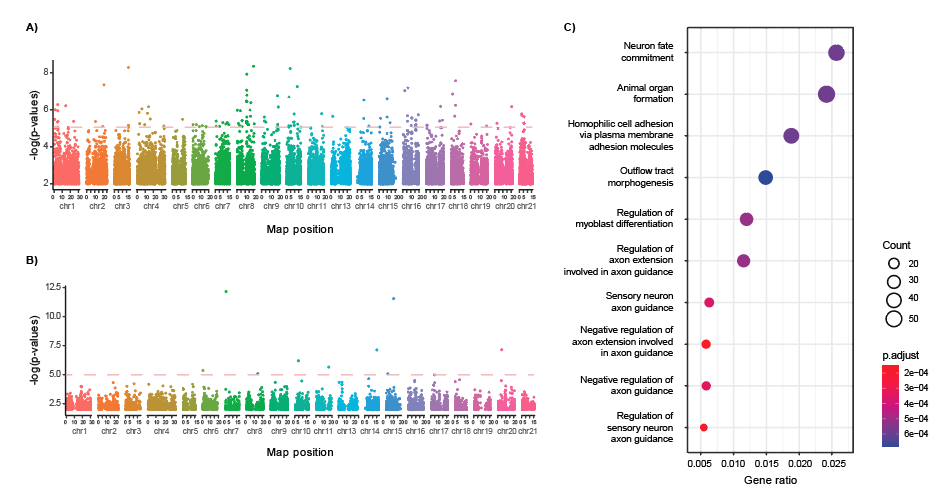

**Supplementary Figure 11.** A potential structural variant found located on a long non-coding RNA (*lncrna*) in the hypermutable pedigree *fin_ryt22*. A deletion in the region of chr10:4,393,834-4,393,920 in both parents was found, which was inherited by all of their offspring. The first two lanes are the mother and the father in *fin_ryt22*, whereas in the following lane of vcf genotypes, light blue, dark blue and white represents deletion homozygote, deletion heterozygote, and the homozygote of the ancestral states.

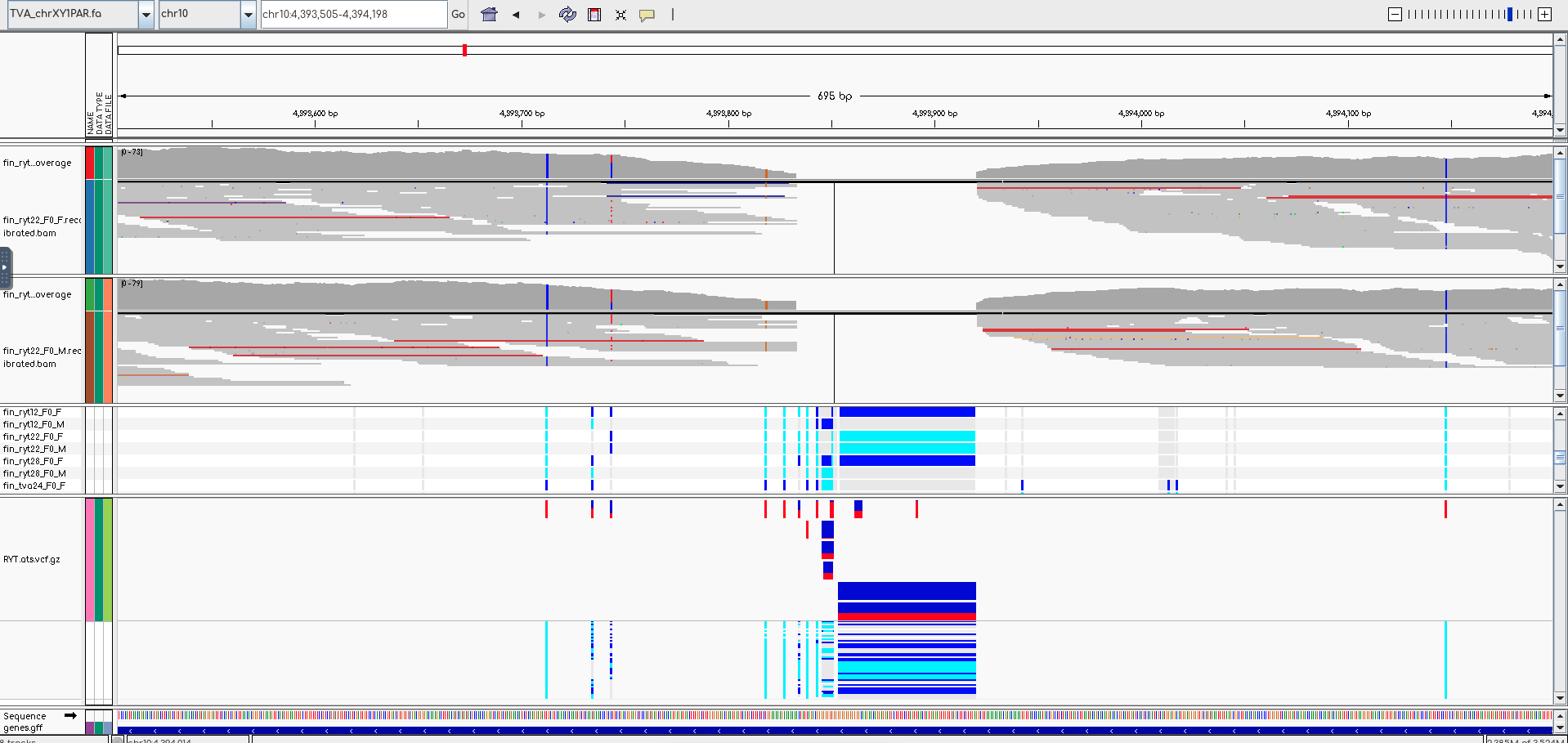

**Supplementary Figure 12.** Illustration of the *de novo* mutation occurring at different chromosomes and the filtering setting based on the ploidy.

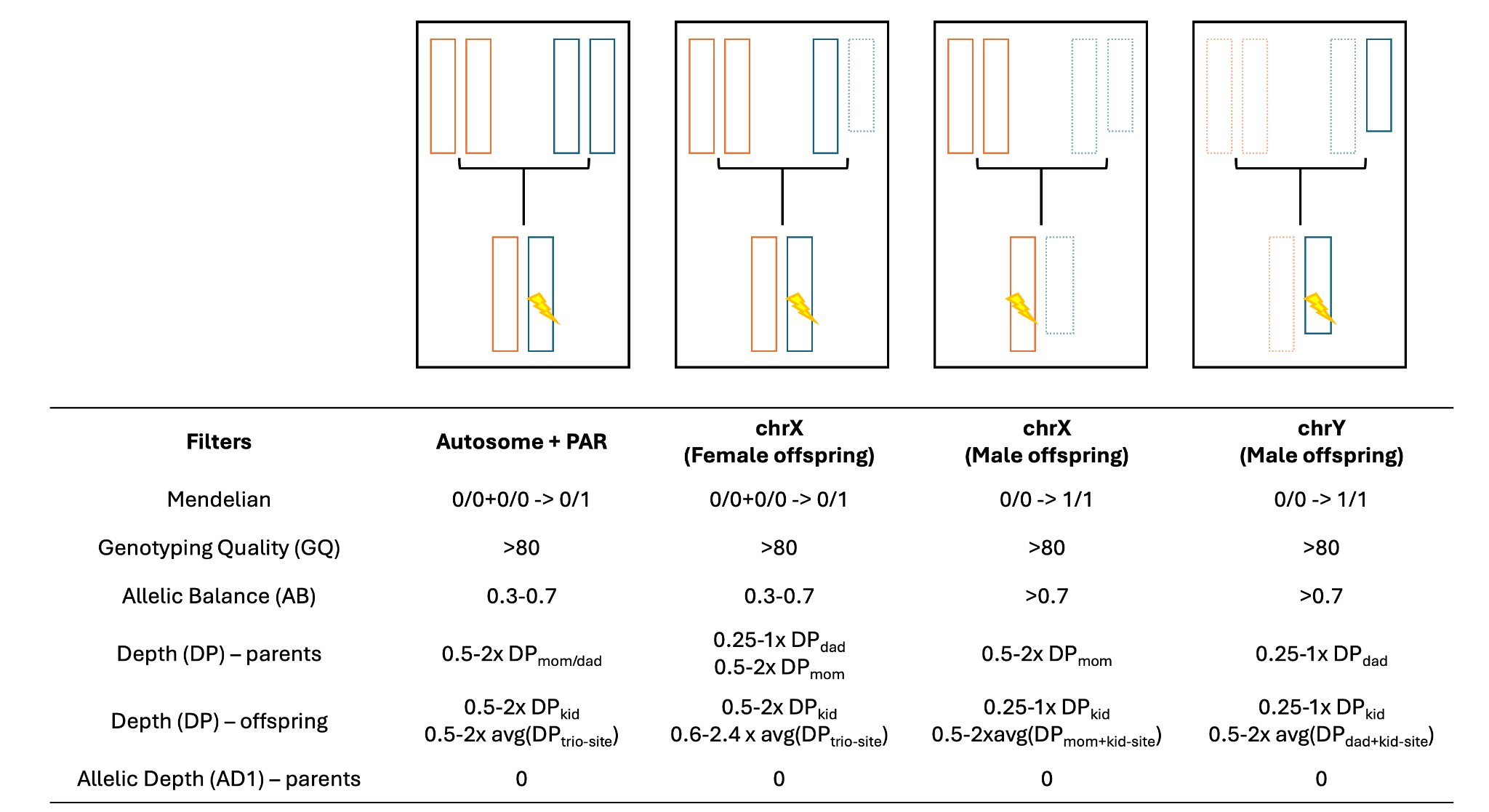

**Supplementary Figure 13.** DNM verification examples for A) IGVtools plot and B) PCR results for a true mutation chr4:28263805 identified in swe_ume34_F1F01 (or LG4:27532124 with mapping to Version 7 assembly). The three panels for the both plots were sequentially from the mother (swe_ume34_F0F01), father (swe_ume34_F0M01) and the offspring.

A) IGV plot

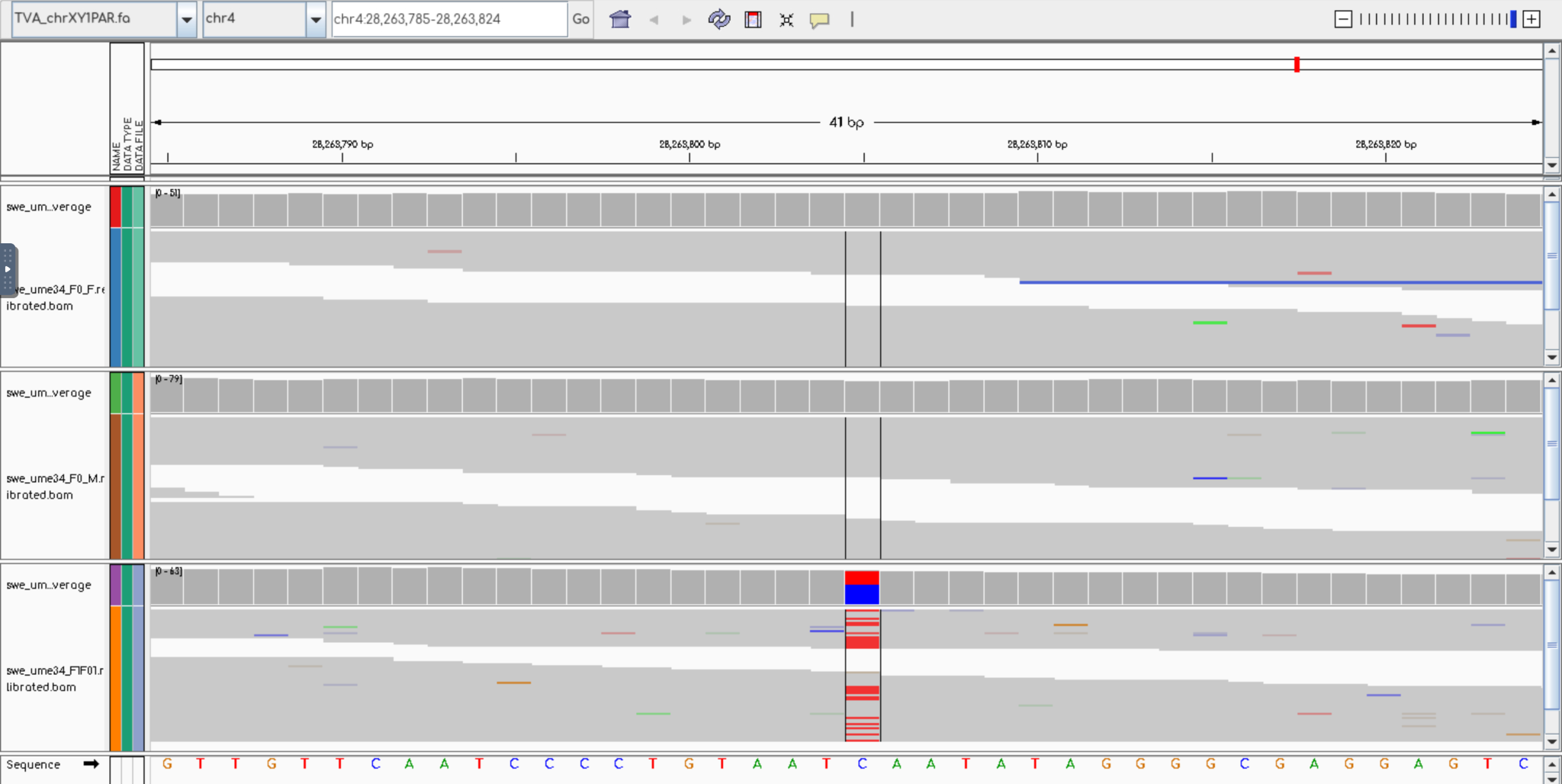

B) Sanger sequencing results shown in Geneious (2025.v.3)

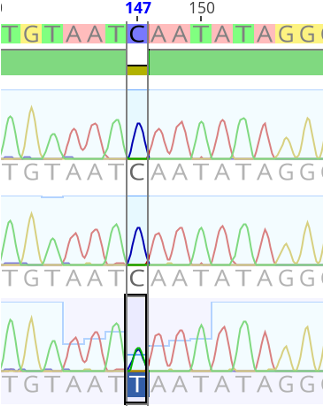
